# Stress-induced nucleoid remodeling in *Deinococcus radiodurans* is associated with major changes in HU abundance and dynamics

**DOI:** 10.1101/2023.10.18.562934

**Authors:** Pierre Vauclare, Jip Wulffelé, Françoise Lacroix, Pascale Servant, Fabrice Confalonieri, Jean-Philippe Kleman, Dominique Bourgeois, Joanna Timmins

**Author notes:** These authors contributed equally to this work (Pierre Vauclare, Jip Wulffelé).

## Abstract

Bacteria have developed a wide range of strategies to respond to stress, one of which is the rapid large-scale reorganization of their nucleoid, which is often associated with a major reprogramming of the gene expression profile. Nucleoid associated proteins (NAPs) are believed to be major actors in this process, but the molecular mechanisms underlying stress-induced nucleoid remodeling remain poorly understood. Here, using the radiation resistant bacterium, *D. radiodurans*, as a model, and advanced fluorescence microscopy approaches, we examined the changes in nucleoid morphology and compaction induced by either entry into stationary phase or exposure to UV-C light, and characterized the associated changes in abundance and dynamics of the major NAP in *D. radiodurans*, the heat-unstable (HU) protein. While both types of stress induced a similar macroscopic rearrangement of the nucleoid into a more compact structure, HU diffusion was significantly reduced in stationary phase cells, but was instead dramatically increased following exposure to UV-C, suggesting that the underlying mechanisms of remodeling are distinct. Furthermore, a detailed comparison of the cellular response to sublethal and lethal doses of UV-C light revealed that UV-induced nucleoid remodeling involves a rapid nucleoid condensation step associated with increased HU diffusion and abundance, followed by a slower decompaction phase to restore normal nucleoid morphology and HU dynamics, before cell growth and division can resume. Together, these findings shed light on the diversity and complexity of stressed-induced nucleoid remodeling processes in bacteria.

## INTRODUCTION

An organism’s capacity to survive depends largely on its ability to sense and respond to changes in its environment. This is particularly true for bacteria that are constantly exposed to adverse environmental conditions, affecting their physiology, growth and survival. To respond to such stress, bacteria have developed a wide range of strategies. Among these, large-scale reorganization of the overall architecture of the genome appears to be one of the most rapid and effective strategies, especially in response to sudden stress (Dame *et al*., 2020; Hołówka and Zakrzewska-Czerwińska, 2020), and is generally associated with temporal changes in the gene expression profile of the bacterium (Steil *et al*., 2005; Kar *et al*., 2005; Mitosch *et al*., 2019; Remesh *et al*., 2020). Stress-induced nucleoid remodeling is a widespread mechanism shared across many bacterial species including human pathogens and often results in a more compact nucleoid. It typically occurs in bacteria entering stationary phase or undergoing sporulation (Meyer and Grainger, 2013; Floc’h *et al*., 2019; Dame *et al*., 2020; Szafran *et al*., 2020), states in which bacteria can withstand major changes in their environment, such as reduced nutrient availability (Piggot and Hilbert, 2004; Hengge, 2011). Nucleoid remodeling has also been reported in bacteria exposed to antibiotics, low pH, or oxidative stress (Chawla *et al*., 2018; Morikawa *et al*., 2019; Hołówka and Zakrzewska-Czerwińska, 2020; Remesh *et al*., 2020). Nucleoid condensation has been proposed to insulate genomic DNA from its environment by creating a physical barrier between the genome and the stress factors (Shechter *et al*., 2013; Hołówka and Zakrzewska-Czerwińska, 2020), and to facilitate DNA repair by limiting the diffusion of broken chromosomes (Zimmerman and Battista, 2005). Despite the well-recognized importance of nucleoid remodeling as a major response to stress, the molecular mechanisms underlying this reorganization remain poorly understood.

The bacterial nucleoid displays a highly hierarchical organization, composed of macro- and micro- domains that play key roles in genome maintenance and regulation of gene expression (Dame *et al*., 2020; Szafran *et al*., 2020). This packaging of the genome is achieved by several factors, including DNA supercoiling, molecular crowding and the abundant nucleoid-associated proteins (NAPs). NAPs are small positively charged proteins that interact with the genomic DNA to define the architecture of the nucleoid. NAPs preferentially bind to DNA in a non-specific manner and control DNA packaging through DNA bending, wrapping or bridging distant strands. However, they have also been reported to bind specifically to DNA motifs thereby contributing to regulating processes such as gene-specific transcription, replication, recombination and repair (Dillon and Dorman, 2010; Verma *et al*., 2019; Dame *et al*., 2020). A large number of genes involved in bacterial viability, metabolism and stress response are thus directly or indirectly regulated by NAPs (Azam and Ishihama, 1999; Macvanin and Adhya, 2012; Remesh *et al*., 2020). Environmental changes have been shown to affect the expression level and/or DNA binding properties of NAPs, notably through post-translational modifications such as phosphorylation, acetylation, methylation, succinylation, oxidation and deamination, in turn affecting the DNA topological organization (Gupta *et al*., 2014; Ghosh *et al*., 2016; Okanishi *et al*., 2017; Dilweg and Dame, 2018). In many bacteria, NAPs and in particular Dps (DNA-binding protein from starved cells) proteins have been shown to contribute to nucleoid condensation and DNA protection, notably in stationary phase (Nair and Finkel, 2004; Haikarainen and Papageorgiou, 2010; Sato *et al*., 2013; Karas *et al*., 2015). NAPs are thus likely to be key players in stress-induced nucleoid remodeling (Boor, 2006; Boutte and Crosson, 2013).

Most of our current knowledge on nucleoid organization comes from the study of rod-shaped bacteria like *E. coli*, while studies on cocci are still sparse. A fascinating model is the well-known *Deinococcus radiodurans,* a non-pathogenic gram-positive coccus (Makarova *et al*., 2001; Cox and Battista, 2005). The exceptional resistance of *D. radiodurans* to a wide range of genotoxic stresses including desiccation, oxidizing agents, and ionizing and UV radiation, has been attributed to its effective anti-oxidant and reactive oxygen species (ROS)-scavenging strategies combined with a highly efficient DNA repair machinery (Krisko and Radman, 2013; Timmins and Moe, 2016). *D. radiodurans* possesses a complex multipartite and multicopy genome arranged into a condensed nucleoid (Zimmerman and Battista, 2005; Floc’h *et al*., 2019). Interestingly, it was shown that *D. radiodurans* nucleoids remain nonetheless highly dynamic, capable of adopting multiple distinct structures in exponential phase as cells progress through their cell cycle (Floc’h *et al*., 2019). Strikingly, in comparison with radiosensitive bacteria (Azam and Ishihama, 1999), the high level of chromosome compaction in *D. radiodurans* is managed by a small number of versatile NAPs (Bouthier de la Tour *et al*., 2013; Chen *et al*., 2020). Indeed, amongst the five most abundant NAPs found in *E. coli* (HU, IHF, H-NS, Fis and Dps), only two of them, HU and two Dps variants (Dps1 and Dps2), are present in *D. radiodurans* (Makarova *et al*., 2001; Romao *et al*., 2006; Cuypers *et al*., 2007). These two Dps proteins, however, do not appear to play a major role in nucleoid organization and condensation as observed in *E. coli*, but have instead been shown to contribute to protection against oxidative stress by regulating the availability of manganese and iron in the cell (Nguyen *et al*., 2009; Santos *et al*., 2015; Santos *et al*., 2019).

The ubiquitous histone-like HU protein is the most widespread and abundant NAP in bacteria (Stojkova *et al*., 2019). Extensive studies have shown that HU is a dimeric protein that exists mostly as a homodimer, but also as a heterodimer in enterobacteria such as *E. coli*, where it can form either homo-(HUα-HUα) or heterodimers (HUα/Huβ) (Nguyen *et al*., 2009; Grove, 2011; Hammel *et al*., 2016). Its role in nucleoid compaction appears multifold. Recent single-particle tracking experiments have suggested that HU mainly contributes to nucleoid organization via transient non-specific interactions (Bettridge *et al*., 2021; Stracy *et al*., 2021). At the same time, *in vitro* studies have shown that HU exhibits both non-specific binding to unstructured DNA and higher affinity specific binding to distorted DNA structures (Kamashev and Rouviere-Yaniv, 2000; Ghosh and Grove, 2004; Verma *et al*., 2023). HU is also able to multimerize on DNA, leading to DNA stiffening or compaction depending on its concentration (Hammel *et al*., 2016; Remesh *et al*., 2020; Chen *et al*., 2020). HU-DNA interactions have indeed been proposed to be affected by the local HU concentration, but also by post-translational modifications, in particular phosphorylation and acetylation, which contribute to the fine tuning of the global architecture of the nucleoid (Gupta *et al*., 2014; Ghosh *et al*., 2016; Dilweg and Dame, 2018; Hou *et al*., 2022). Beyond its role in nucleoid organization, HU, like other NAPs, has been reported to play a role in transcriptional regulation, DNA replication and DNA repair (Aki, 1997; Oberto *et al*., 2009; Hammel *et al*., 2016; Li and Waters, 1998; Kamashev and Rouviere-Yaniv, 2000). HU-deficient *E. coli* cells remain viable with a modest change in phenotype, but have been found to be more sensitive to UV irradiation (Huisman *et al*., 1989), suggesting a role for HU in bacterial stress-response.

In *D. radiodurans*, unlike in *E. coli*, the *hbs* gene encoding HU is essential for cell viability, and the progressive depletion of *D. radiodurans* HU (DrHU) using a thermosensitive plasmid leads to nucleoid decondensation, fractionation and eventually cell lysis (Nguyen *et al*., 2009). Together with the DNA gyrase that modulates the topology of the genomic DNA, DrHU thus appears to be the primary structuring protein orchestrating nucleoid organization in *D. radiodurans* (Bouthier De La Tour *et al*., 2015). Interestingly, DrHU harbors an additional 30 amino acid long N-terminal lysine-rich extension, that is reminiscent of the C-terminal domain of eukaryotic histone H1, which is also involved in DNA compaction (Ghosh and Grove, 2006; Bouthier De La Tour *et al*., 2015).

In this work, using notably a combination of 3D spinning-disk microscopy and single-particle tracking experiments, we have performed an in-depth study of nucleoid remodeling in *D. radiodurans* and its associated changes in HU abundance and dynamics, induced by two types of stress: (i) entry into stationary phase, a stress that occurs frequently in the natural environment, during which bacteria are faced with resource-limited conditions preventing any further growth (Gefen *et al*., 2014; Dworkin and Harwood, 2022), and (ii) exposure to intense UV-C light that produces extensive DNA damage, and notably the formation of pyrimidine photoproducts, but also ROS (Richa *et al*., 2015). The data reveal that the nucleoid adopts a compacted and spherical morphology both in stationary phase and immediately after exposure to UV-C radiation. However, strikingly, single-particle tracking of HU revealed that while HU diffusion decreases during stationary phase, it dramatically increases following exposure to UV-C, suggesting that the underlying mechanisms are distinct. Furthermore, by comparing the cellular response of *D. radiodurans* to sublethal and lethal doses of UV-C light, we determined that the nucleoid remodeling process starts with a rapid nucleoid condensation step associated with increased HU abundance and diffusion, which is then followed by a slower decompaction phase, seen only in surviving cells, during which genome integrity and nucleoid organization are restored before DNA replication and cell growth can resume. Together, these findings highlight the key role of HU in stress-induced nucleoid remodelling, but also the complexity of this phenomenon in bacteria.

## RESULTS

### Entry into stationary phase induces nucleoid compaction associated with decreased HU dynamics

In our previous work, we characterized the size and morphology of *D. radiodurans* nucleoids, revealing a great diversity of nucleoid shapes in exponential growth phase and more condensed and homogeneous nucleoids in stationary phase (Floc’h et al., 2019). We confirmed these findings by 3D confocal imaging of membrane (Nile Red)- and DNA (Syto9)-stained wild-type D. radiodurans (DR^WT^) cells and of a genetically engineered strain of *D. radiodurans* expressing HU-mCherry from its endogenous locus (DR^HUmCh^) (SI Fig. S1A). Imaging was performed on bacteria in both exponential and early stationary growth phases (Fig. 1A and SI Fig. S2A-B). In DR^HUmCh^, HU-mCherry covers the genomic DNA and serves as a proxy to follow changes in the organization of the nucleoid (SI Fig. S1A). As reported previously, the mean volume of DR^HUmCh^ nucleoids is slightly higher than that of DR^WT^, possibly as a result of steric hindrance caused by the mCherry label (Floc’h *et al*., 2019). For both strains, we evaluated the fraction of rounded nucleoids, and determined the volume and sphericity of individually segmented nucleoids, so as to perform a thorough statistical comparison of exponential versus stationary phase cells. We found that in stationary phase, almost 90% of cells exhibited a rounded nucleoid (SI Fig. S2B) and the mean nucleoid volume was decreased by almost 50% compared to that in exponential phase, going from 1.01 ± 0.34 µm^3^ to 0.53 ± 0.19 µm^3^ (Fig. 1B). The sphericity, in contrast, increased significantly (+12%) and displayed a markedly reduced distribution in stationary phase (Fig. 1B), confirming that the nucleoid morphology is indeed much more homogeneous in this growth phase. Similar observations were made on the DR^HUmCh^ strain with a 40% drop in the nucleoid volume and a significant increase in the sphericity (Fig. 1B). Western blot detection of HU-mCherry in cell extracts of DR^HUmCh^ revealed that HU levels were significantly higher (3-fold) in stationary phase cells compared to exponential phase cells (Fig. 1C and SI Fig. S2C).

**Figure 1:**
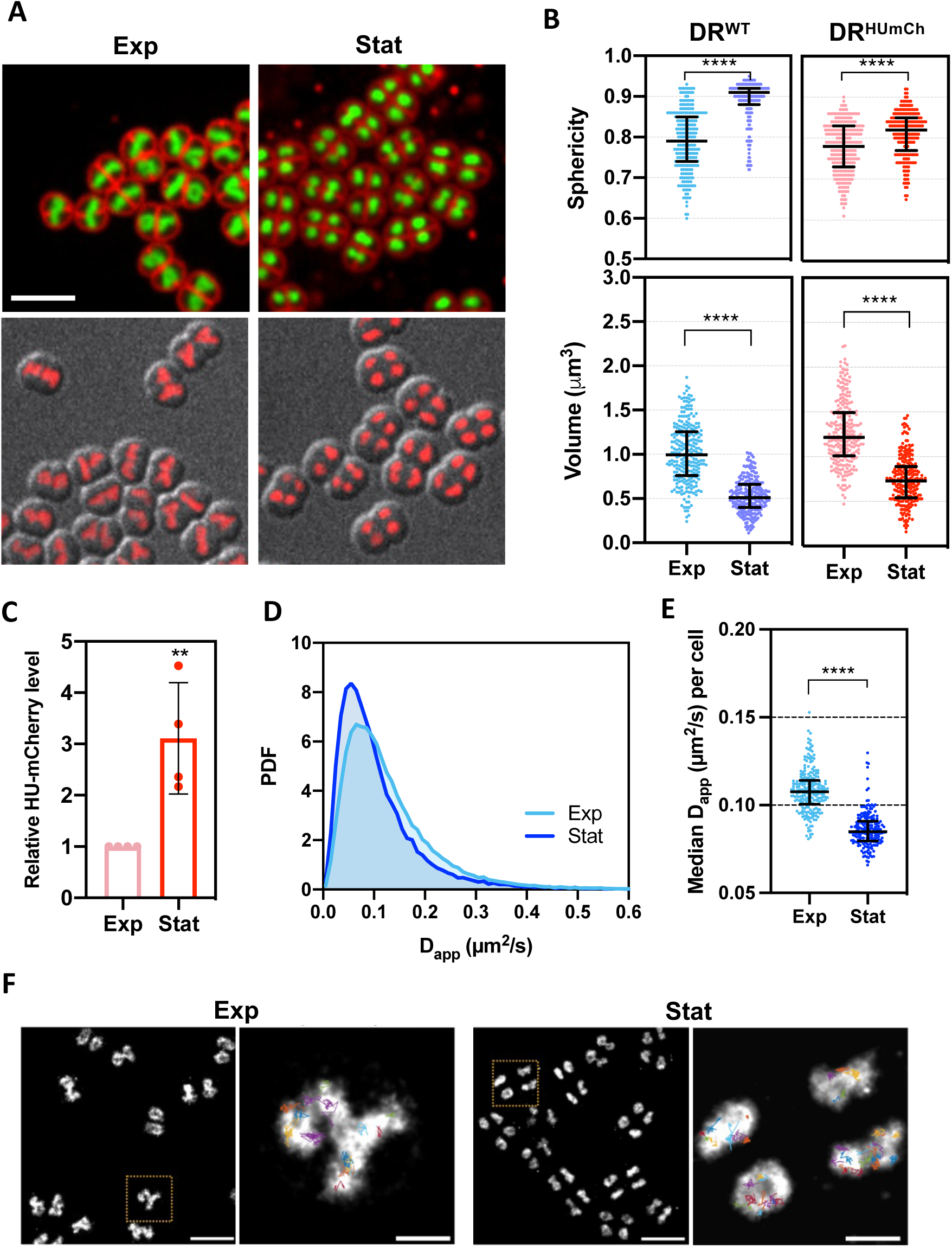
Growth phase-dependent nucleoid remodeling and associated changes in HU dynamics. (A) Representative images of DR^WT^ cells (upper panels) stained with Syto 9 and Nile Red and DR^HUmCh^ (lower panels) in exponential (left) and stationary (right) phase. Scale bar: 5 µm. (B) Nucleoid volume and sphericity of DR^WT^ (blue) and DR^HUmCh^ (red) strains in exponential (light color) and stationary phase (dark color) cells (n=250). Error bars represent the median and interquartile range. **** p<0.0001, Kruskal-Wallis statistical test performed in GraphPad Prism 8. (C) Expression levels of HU-mCherry in exponential and stationary phase (n=3). (D) Histogram distribution of the apparent diffusion coefficient (D_app_) of HU-mEos4b in exponential (light blue; n=231) and stationary (dark blue; n=196) phase DR^HUmEos^ cells. PDF: Probability density function. (E) Median D_app_ of HU-mEos4b per cell in exponential (light blue; n=231) and stationary phase (dark blue; n=196) cells. Error bars represent the median and interquartile range. **** p<0.0001, Kruskal-Wallis statistical test performed in GraphPad Prism 8. (F) Representative images of exponential (left) and stationary (right) phase DR^HU-mEos^ nucleoids. The right panel is a close up view of the boxed cells in the left panel and individual HU-mEos4b tracks are overlayed on the epi-fluorescence image. Scale bar upper panel: 0.5 µm. Scale bar lower panel: 3 µm.

Next, we compared the dynamics of HU in exponential and stationary phase cells by single-particle tracking (spt; SI Fig. S3A). For this, we constructed a *D. radiodurans* strain expressing HU fused to mEos4b (De Zitter *et al*., 2019), DR^HUmEos^ (SI Fig. S1A), enabling an in-depth characterization of HU dynamics (Fig. 1D-F). PALM imaging of this strain revealed the same variety of nucleoid shapes as seen by confocal microscopy in exponential phase and a similar change in nucleoid morphology upon entering stationary phase (Fig. 1F). We plotted the apparent diffusion coefficient (D_app_) values calculated from the mean square jump distance of 4 displacements per track in exponential and stationary phase cells (Fig. 1D), and extracted the median D_app_ per cell (Fig. 1E). We observed that nucleoid rounding and compaction upon entry into stationary phase was accompanied by a marked reduction in the dynamics of HU, with an almost 20% lower D_app_ than that observed in exponential phase cells, suggesting a major reorganization of the HU-DNA assembly in these highly condensed nucleoids. We chose not to fit the spt data to distinct populations of molecules as reported in several earlier studies (Kapanidis *et al*., 2018; Bettridge *et al*., 2021; Stracy *et al*., 2021), since simulations revealed that such an analysis would likely be biased by the strong and heterogeneous confinement encountered by HU molecules in the various nucleoid morphologies observed in *D. radiodurans.*A detailed discussion of this issue is provided in the supplementary data (SI discussion).

### UV-C light elicits a rapid dose-dependent compaction of the nucleoid

*D. radiodurans* is known to exhibit an outstanding resistance to UV light (Harsojo *et al*., 1981; Battista, 1997; Slade and Radman, 2011). We exposed exponential and stationary phase DR^WT^ cultures to a range of UV-C doses. Up to 1.9 kJ/m^2^, the survival of *D. radiodurans* was largely unaffected, but rapidly dropped beyond this dose, regardless of whether cells were in exponential or stationary phase (Fig. 2A). In contrast, survival of *E. coli* under the same experimental conditions was severely impacted already at a dose of 0.2 kJ/m^2^. The UV resistance of DR^HUmCh^ and DR^HUmEos^ strains was similar to that of DR^WT^, with between 30 and 50% cell survival at the sublethal dose of 1.9 kJ/m^2^ and less than 0.01% survival at the lethal dose of 12.0 kJ/m^2^ (SI Fig. S1B).

**Figure 2:**
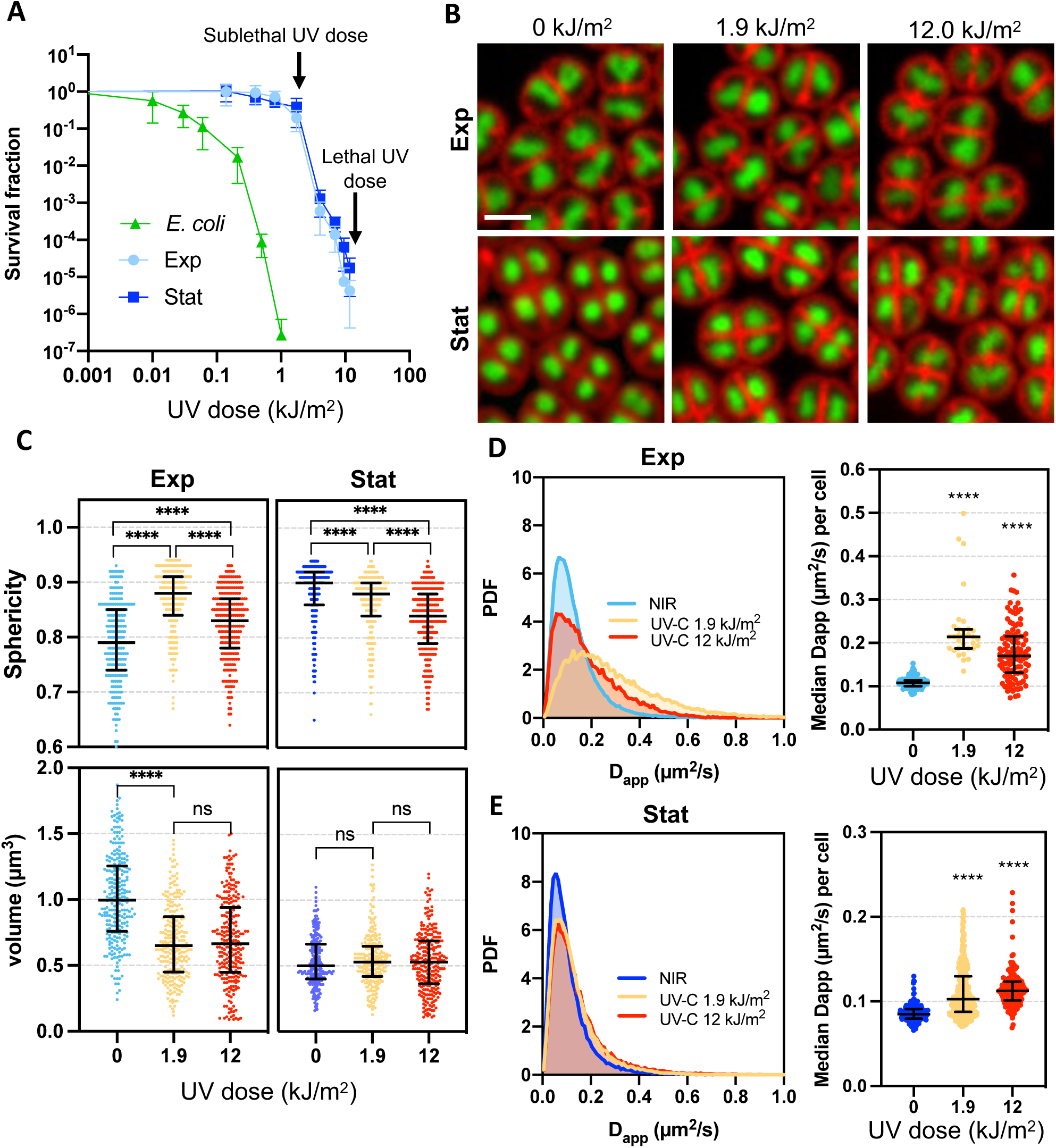
Dose-dependent effects of UV-C light on nucleoid organization. (A) Survival curves of exponential (light blue) and stationary (dark blue) DR^WT^ and *E. coli* (green) at different UV doses. The sublethal (1.9 kJ/m^2^) and lethal (12.0 kJ/m^2^) doses used for the subsequent studies are indicated with arrows. Data represent the mean and standard deviation of at least 3 independent experiments. (B) Representative images of exponential (top panel) and stationary phase (lower panel) DR^WT^ cells stained with Syto9 and Nile Red before (0 kJ/m^2^; left) and after exposure to 1.9 (middle) or 12.0 kJ/m^2^ UV-C light. Scale bar: 2.5 µm. (C) Nucleoid volume and sphericity of exponential (left) and stationary (right) DR^WT^ cells before (0 kJ/m^2^; blue) and after exposure to 1.9 (orange) or 12.0 kJ/m^2^ (red) UV-C light cells (n=250). Error bars represent the median and interquartile range. ns: non-significant, **** p<0.0001, Kruskal-Wallis statistical test performed in GraphPad Prism 8. (D-E) Histogram distribution of the apparent diffusion coefficient (D_app_) of HU-mEos4b (left) and median D_app_ of HU-mEos4b per cell (right) in exponential (top panel) and stationary (lower panel) DR^HUmEos^ before (blue; n=231 for exp and 196 for stat) and after exposure to 1.9 (orange; n=32 for exp and 356 for stat) or 12.0 kJ/m^2^ (red; n=114 for exp and 152 for stat) UV-C light. PDF: Probability density function. Error bars represent the median and interquartile range. **** p<0.0001, Kruskal-Wallis statistical test performed in GraphPad Prism 8.

UV-C-induced changes in nucleoid volume and morphology were then evaluated by 3D spinning-disk microscopy, as described above, after irradiation of *D. radiodurans*. Due to the time needed for sample preparation, confocal images were typically acquired 30 minutes after irradiation (cf. Materials and Methods). A dose-dependent compaction of the nucleoid of exponentially growing DR^WT^ and DR^HUmCh^ was observed up to 1.9 kJ/m^2^ (Fig. 2B-C and SI Fig. S4 and S5) reaching a minimal volume of 0.65 µm^3^ for DR^WT^ and 1.11 µm^3^ for DR^HUmCh^, corresponding to respectively a 35% and 18% decrease in the mean nucleoid volume. As in the case of stationary phase cells, this nucleoid compaction was accompanied by an increased sphericity and a marked reduction in the spread of the sphericity values, suggesting the formation of more compact and homogeneous nucleoids in the cell population (Fig. 2B-C and SI Fig. S4 and S5). Of note, in DR^WT^, beyond the sublethal dose of 1.9 kJ/m^2^, the nucleoid volume remained constant at its minimal value, whereas the mean sphericity seemed less affected by the higher UV-C doses, suggesting that nucleoid rounding might be partly impaired at these highly cytotoxic UV-C doses (Fig. 2B-C and SI Fig. S4 and S5). The same experiments were then performed on stationary DR^WT^ and DR^HUmCh^ cultures. While no further nucleoid compaction was observed for DR^WT^ stationary phase nucleoids following exposure to UV-C light (Fig. 2B-C and SI Fig. S4A and S5A), the volume of stationary phase DR^HUmCh^ nucleoids did show a significant dose-dependent reduction going from 0.74 µm^3^ in non-irradiated samples to 0.39 µm^3^ in response to 12.0 kJ/m^2^ (SI Fig. S4B and S5B). This difference may be explained by the different levels of compaction of stationary phase nucleoids in the two strains before irradiation.

Having established the global changes in nucleoid shape and size following exposure to UV-C light, we next examined the effects of sublethal (1.9 kJ/m^2^) and lethal (12.0 kJ/m^2^) doses of UV-C radiation on HU dynamics using the DR^HUmEos^ strain. We first ensured that UV-C illumination does not change the fluorescence properties of mEos4b (SI Fig. S1A and Fig. S3B). We then observed that, although UV-C induced a clear compaction of the nucleoid as in cells entering stationary phase (SI Fig. S6), UV-C also led to a significant increase in the apparent diffusion coefficient (D_app_) of HU in irradiated cells (at both sublethal and lethal doses) in both exponentially and stationary phase cells (Fig. 2D-E). In exponential cells, the median D_app_ of HU-mEos4b in irradiated cells was approximately twice that measured in non-irradiated samples, while in stationary phase cells, HU diffusion was ∼30% higher after irradiation. The more compact structure of stationary phase nucleoids may be more restrictive for HU diffusion.

To determine whether the increased HU diffusion coefficient observed after exposure to UV-C light could be due to increased mobility of the genomic DNA as a result of UV-induced DNA damage, we probed the dynamics of the *OriC* and *Ter* loci of chromosome 1 (the largest chromosome of *D. radiodurans*, corresponding to ∼80% of the genome) by spt using a heterologous *parS*-ParB system (SI Fig. S7A-B) previously used to track these loci in *D. radiodurans* cells (Passot *et al*., 2015; Floc’h *et al*., 2019). Exponentially growing *D. radiodurans* strains in which either *OriC* or *Ter* were labelled with ParB-GFP were thus irradiated with a sublethal dose of UV-C light and imaged after 1 hour of recovery. No noticeable changes in the mobility of chromosome 1 loci were observed after exposure to UV-C light (SI Fig. S7C-D), suggesting that changes in HU dynamics probably reflect altered HU-DNA interactions and not increased mobility of the damaged DNA.

### Cell and nucleoid recovery after a sublethal dose of UV-C light involves three distinct stages

To decipher the different stages of UV-C-induced nucleoid remodeling, we carried out a detailed analysis of the response of exponentially growing *D. radiodurans* cells to a sublethal dose of UV-C light up to 24h following irradiation (Fig. 3 and 4, and SI Fig. S8 and S9). For this purpose, irradiated cells (DR^WT^, DR^HUmCh^ and DR^HUmEos^) were transferred back into fresh medium immediately after irradiation and samples were then collected at different timepoints and subjected to a variety of measurements: (i) optical density measurements to determine growth curves after irradiation, (ii) 3D confocal microscopy on DR^WT^ and DR^HUmCh^ to assess cell growth and division, changes in size and morphology of the nucleoids, (iii) western blot analysis of DR^HUmCh^ cell extracts and 3D confocal microscopy on DR^HUmCh^ to follow the abundance of HU, and (iv) spt experiments on DR^HUmEos^ cells to probe the dynamics of HU. Before carrying out these analyses, we confirmed that the three strains of *D. radiodurans* respond in a similar way to exposure to UV-C light by establishing their growth curves and determining the size and morphology of their nucleoids at selected timepoints after irradiation (SI Fig. S8).

**Figure 3:**
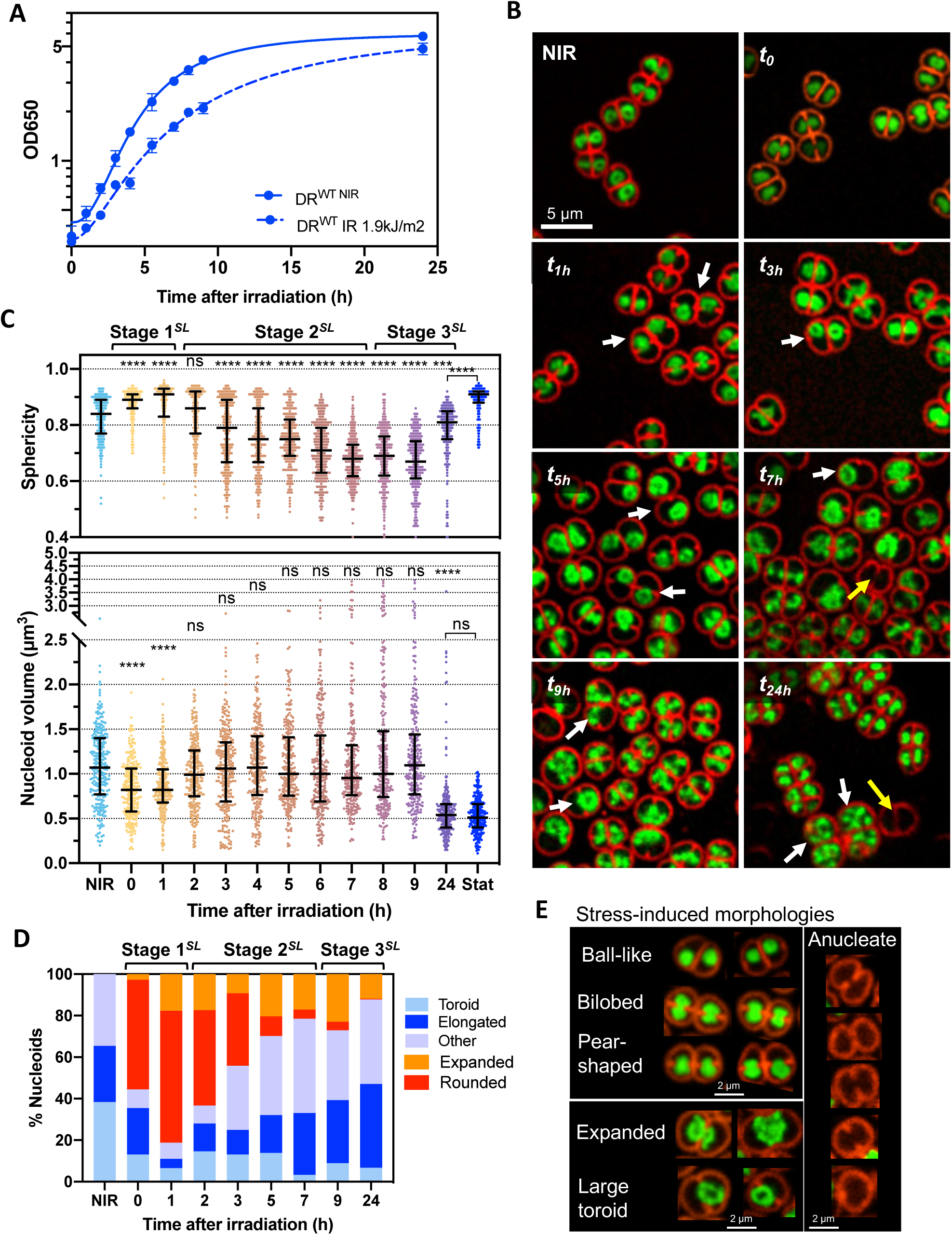
Effects of sublethal UV-C exposure on nucleoid organization and morphology. (A) Growth curves of non-irradiated (NIR; full line) and irradiated DR^WT^ (1.9 kJ/m^2^; dashed line). Data represent the mean and standard deviation of at least 3 independent experiments. (B) Representative images of DR^WT^ cells stained with Syto9 and Nile Red at different timepoints after exposure to 1.9 kJ/m^2^ UV-C light. White and yellow arrows indicate cells with off-center nucleoids and anucleate cells respectively. Scale bar: 5 µm. (C) Nucleoid volume and sphericity of DR^WT^ cells at different timepoints after exposure to 1.9 kJ/m^2^ UV-C light (n=250). Error bars represent the median and interquartile range. ns: non-significant, **** p<0.0001, Kruskal-Wallis statistical test performed in GraphPad Prism 8. (D) Evolution of nucleoid morphologies during the recovery from exposure to 1.9 kJ/m^2^ UV-C light (n>450). (C)-(D) The three different stages of the recovery phase are indicated above the plots. (E) Examples of the most common UV-C-induced nucleoid and cell morphologies observed by confocal microscopy of Syto9 and Nile Red stained DRWT cells before and after exposure to 1.9 kJ/m^2^ UV-C light. In (D), rounded nucleoids correspond to ball-like, bilobed and pear-shaped nucleoids, while expanded nucleoids refer to both expanded and large toroids as depicted in (E).

**Figure 4:**
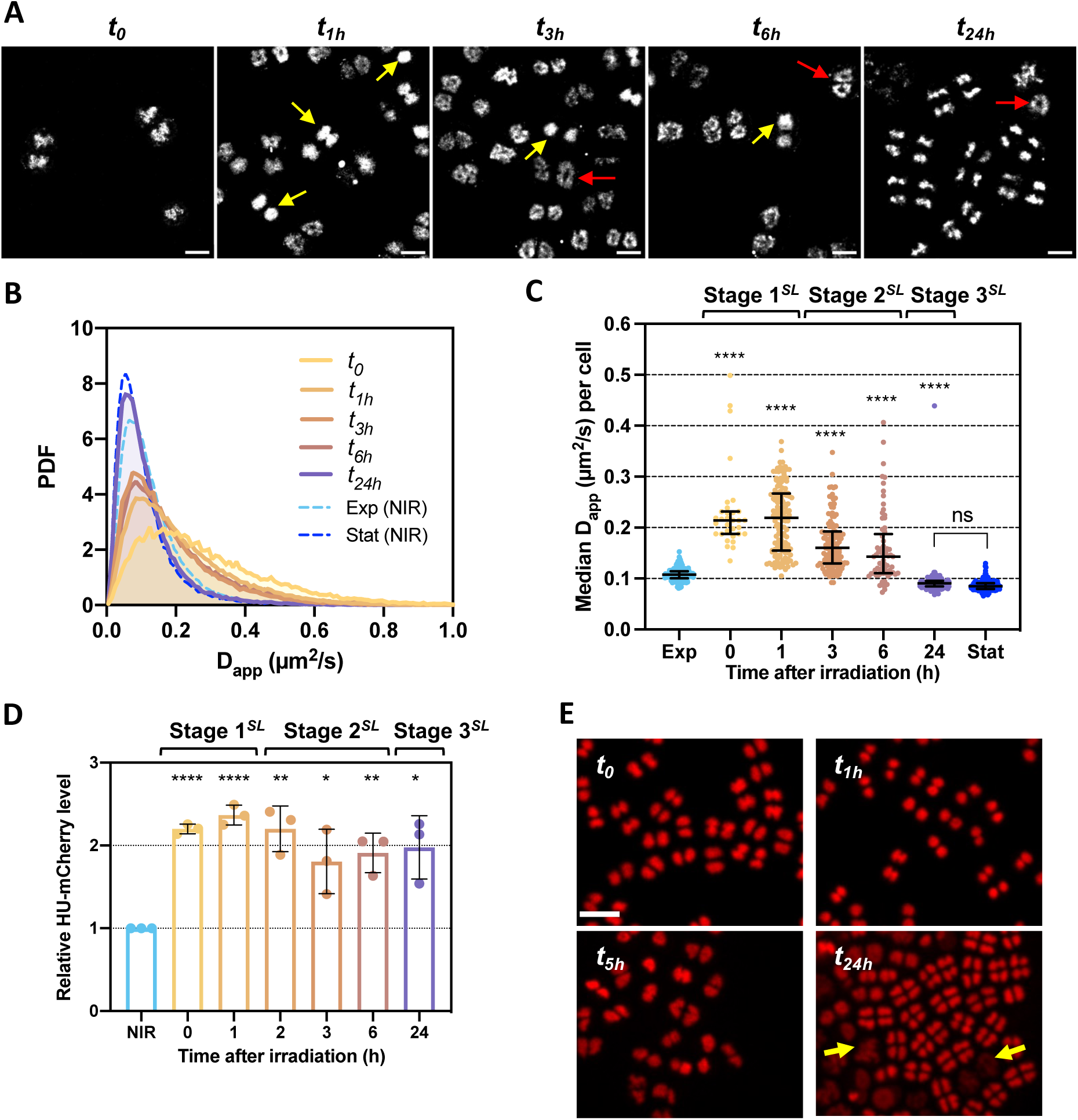
Effects of sublethal UV-C exposure on the dynamics and abundance of HU. (A) Representative images of DR^HUmEos^ nucleoids at different timepoints after exposure to 1.9 kJ/m^2^ UV-C light. Yellow and red arrows indicate rounded and expanded nucleoids respectively. Scale bar: 2 µm. (B) Histogram distribution of the apparent diffusion coefficient (D_app_) of HU-mEos4b in non-irradiated stationary DR^HUmEos^ (dashed blue line) cells and at different timepoints after exposure to 1.9 kJ/m^2^ UV-C light (orange to purple; n>30). PDF: Probability density function. (C) Median D_app_ of HU-mEos4b per cell in non-irradiated exponential (light blue) and stationary (dark blue) DR^HUmEos^ cells and at different timepoints after exposure to 1.9 kJ/m^2^ UV-C light (orange to purple; n>30). Error bars represent the median and interquartile range. ns: non-significant, **** p<0.0001, Kruskal-Wallis statistical test performed in GraphPad Prism 8. (D) Relative HU-mCherry levels determined by Western blot analysis at different timepoints after exposure to 1.9 kJ/m^2^ UV-C light (orange to purple; n=3). The intensity of NIR samples was set to 1. Error bars represent the standard deviation. * p<0.05, ** p<0.01, **** p<0.0001, unpaired t-test performed in GraphPad Prism 8. (C)-(D) The three different stages of the recovery phase are indicated above the plots. (E) Representative images of DR^HUmCh^ at different timepoints after exposure to 1.9 kJ/m^2^ UV-C light. Yellow arrows indicate cells with diffuse HU-mCherry staining. Scale bar: 5 µm.

The recovery of *D. radiodurans* bacteria exposed to a sublethal (SL) UV-C dose can be divided into three main stages: stage 1*^SL^* (*t*_0_ to *t*_1h_ post-irradiation) during which cell growth and division are arrested and the nucleoids become round and more condensed, stage 2*^SL^* (*t*_1h_ to *t*_7h_ post-irradiation) corresponding to nucleoid decompaction and a progressive recovery of cellular activity, and stage 3*^SL^* (*t*_7h_ to *t*_24h_ post-irradiation) in which most cells have recovered native nucleoid and cell morphologies (Table S1). Optical density measurements performed on DR^WT^, DR^HUmCh^ and DR^HUmEos^ strains exposed to a sublethal dose of UV-C light showed a clear arrest of their cell cycle immediately after irradiation causing a 2 to 3 hour delay in their growth curve (Fig. 3A and SI Fig. S8B), which is in good agreement with the time estimated in earlier studies to allow cells to repair the heavy radiation-induced DNA damage (Zahradka *et al*., 2006). We indeed observed that in DR^WT^ both septation and cell growth were rapidly blocked after irradiation and throughout stage 1*^SL^* of recovery. As a result, transition from phase 3 to phase 4 of the cell cycle, corresponding to the onset of septation, was impaired, leading to an increased fraction of phase 3 cells in the population at the end of stage 1*^SL^* (*t*_1h_; SI Fig. S9A-C). Interestingly, the splitting of tetrads into diads (transition from phase 6 to phase 1; SI Fig. S9A) did not appear to be affected by the irradiation, as evidenced by a significant reduction in the proportion of phase 6 cells during stage 1*^SL^* (SI Fig. S9B). No obvious defects in cell morphology or plasma membrane integrity were seen in this first stage (Fig. 3B).

Arrest of the cell cycle immediately after irradiation was accompanied by a marked reduction in the size of the nucleoid, constituting the first step in the nucleoid remodeling process (Fig. 3B-D for DR^WT^ and SI Fig. S8C for DR^HU-mCh^). The minimal nucleoid volume was reached already at *t_0_* (Fig. 3C and SI Fig. S8C), while the change in nucleoid morphology was found to progress more slowly throughout stage 1*^SL^* to reach its maximum at *t*_1h_ with the highest median sphericity value (0.91) and near 80% of cells exhibiting abnormal nucleoid morphologies, of which 75% were rounded (Fig. 3B-D and SI Fig. S10A). Interestingly, the nucleoids of phase 5 cells (last phase before cytokinesis) responded differently to those of other cells, with almost 50% of phase 5 cells maintaining a structured, elongated nucleoid at *t_1h_* (SI Fig. S10B), suggesting that nucleoid compaction may be less efficient in cells in which chromosome segregation is already well advanced. Nucleoid rounding resulted in different nucleoid morphologies depending on the phase of the cell cycle the cells were in when exposed to UV-C light (Fig. 3E, SI Fig. S10C). A majority of cells in early phases of the cell cycle (phases 1-3) bearing mostly crescent-shaped and ring-shaped nucleoids exhibited ball-like nucleoids after irradiation, whereas cells at later stages of the cell cycle (phases 4-5) in which active chromosome segregation was taking place, exhibited bilobed or pear-shaped nucleoids after irradiation (Fig. 3E and SI Fig. S10C). Moreover, irrespective of their phase of the cell cycle, over 30% of cells exhibited off-center nucleoids at the end of stage 1*^SL^* (Fig. 3B and SI Fig. S10D), in contrast with healthy *D. radiodurans* cells in which the nucleoid is always located in the center of the cell, except immediately after chromosome segregation in phases 5 and 6 (Floc’h *et al*., 2019). Analysis of the HU protein revealed that its dynamics (Fig. 4A-C) and abundance (Fig. 4D and SI Fig. S11A-B) also rapidly increased after irradiation, reaching a maximum at the end of stage 1 (*t_1h_*; Fig. 4A-D).

In stage 2*^SL^* (*t*_1h_ to *t*_7h_ post-irradiation; Table S1), HU dynamics and levels were found to progressively decrease again (Fig. 4C-D). This coincided with the start of the reversal of the nucleoid compaction process. As with compaction, the first step in nucleoid decompaction was the recovery of a ‘normal’ nucleoid volume as early as *t*_2h_ (Fig. 3C and SI Fig. S8C), whereas full recovery of the diverse nucleoid shapes typical of exponentially growing *D. radiodurans* cells took several hours and most of stage 2*^SL^* (Fig. 3D and SI Fig. S10A and D). Nucleoid decompaction was also accompanied by a progressive decrease in the median sphericity of nucleoids and a marked increase in the spread of the sphericity values, reflecting the great diversity of nucleoid shapes observed during the recovery stage, most likely resulting from active DNA repair and cell recovery processes (Fig. 3C and SI Fig. S8C).

Cell growth and division were also largely recovered during stage 2*^SL^* as illustrated by the increased optical density of DR^WT^, DR^HUmCh^ and DR^HUmEos^ cultures (Fig. 3A and SI Fig. S8B) and a restored distribution of cells within the different phases of the cell cycle (SI Fig. S9B). A slow increase in the mean cell size was nonetheless observed between *t_1h_* and *t_3h_*, which may have resulted from a slight delay between the recovery of cell growth and that of cell division (SI Fig. S9B-C). Beyond *t_3h_*, most cells appeared to have recovered their ability to grow and divide and the mean cell size decreased progressively as the cell density increased and the bacterial population moved towards stationary phase. However, we also observed during this period the accumulation of a small fraction (<5%) of anucleate cells (*i.e.* no Syto9 staining; Fig. 3B and E and SI Fig. S10D) that typically also exhibited a damaged cell membrane, and a significant number (∼10% cells) of abnormally large cells in which cell division appeared to be impaired or blocked (Fig. 3B and E and SI Fig. S10E). Many of these enlarged cells also exhibited abnormally large nucleoids that were either spread out throughout the cell and exhibited significantly increased volumes or forming very large rings with a largely unchanged nucleoid volume (Fig. 3C-E and SI Fig. 10F). These cells were likely more severely damaged, causing a delay or even an arrest in the nucleoid remodeling process.

In stage 3*^SL^* of the recovery from a sublethal dose of UV-C light (*t*_7h_ to *t*_24h_ post-irradiation; Table S1), a vast majority of cells had fully recovered and exhibited normal growth and division as evidenced by the rapid increase in the optical densities of the cultures (Fig. 3A and SI Fig. S8B) and the changes in cell phase distribution as the cells shifted from exponential to stationary phase (Fig. 3A-C and SI Fig. S9B). Indeed, at *t*_24h_ post-irradiation, these cultures had reached early stationary phase as seen by the accumulation of phase 6-8 cells (Fig. 3B and SI Fig. S9B). Most cells (80-90%) now exhibited normal, structured and centered nucleoids (Fig. 3D and SI Fig. S10D). After a slight reduction of HU abundance during stage 2*^SL^*, HU levels stabilised in stage 3 at ∼2-fold higher level than in non-irradiated exponential phase samples (Fig. 4D). This increased level may result from the cells reaching stationary phase, which we showed in Fig. 1B is associated with a ∼3-fold increase in HU levels. During stage 3*^SL^*, HU diffusion also continued its recovery to reach values under 0.1 µm^2^/sec as obtained in non-irradiated stationary phase cells (Fig. 4B-C and Fig. 1E). Surprisingly, however, at the end of stage 3*^SL^*, around 10% of cells still exhibited major defects (large cells with impaired septation, expanded nucleoids and high HU diffusion), suggesting that these large, abnormal cells appeared to still arise at late stages of recovery, likely as a result of unrepaired DNA damage leading to cell dysfunctions (Fig. 3B-E, Fig. 4B-C, and SI Fig. S10).

### A lethal dose of UV-C light induces a progressive loss of nucleoid structure and cellular functions

We then subjected our three exponentially growing *D. radiodurans* strains (DR^WT^, DR^HU-mCh^ and DR^HU-mEos^) to a lethal dose of UV-C light and followed the effects of this high dose on the cells and their nucleoids in the hours following irradiation (Fig. 5 and 6 and SI Fig. S12). As with the sublethal dose, all three strains responded in a similar manner (SI Fig. S12A-B) and the recovery could also be divided into three stages, but the nature and duration of these stages differed from those observed after exposure to sublethal UV-C. After a lethal UV-C dose, rapid nucleoid rounding and condensation were also observed in stage 1*^L^* (*t*_0_ to *t*_2h_ post-irradiation; Table S1), but this process lasted longer than after the sublethal dose. Indeed, while a reversal of nucleoid compaction was observed at *t_1h_* (corresponding to the onset of stage 2*^SL^*) after a sublethal dose, here, instead, we noted that after the rapid condensation of the nucleoids immediately after irradiation, the mean nucleoid volume (∼0.75 µm^3^) then remained constant for the rest of stage 1*^L^* until *t_2h_* (Fig. 5B-C and SI Fig. S12A and Fig. S13A). The change in nucleoid morphology was also more progressive with the highest sphericity (>0.9) reached at *t_2h_* (Fig. 5B-C). With this higher dose of UV-C, 100% of cells exhibited abnormal nucleoids with ∼95% of cells exhibiting rounded nucleoids and 50% of cells having off-center nucleoids at *t_2h_* (Fig. 5B-D and SI Fig. S13B). These values are considerably higher than those determined for cells exposed to the sublethal dose (Fig. 3D and SI Fig. S10D). During stage 1*^L^*, the levels of HU-mCherry were found to increase immediately after irradiation (*t_0_*) as in response to a sublethal dose of UV-C (Fig. 6A-B) and HU-mEos4b dynamics were also strongly increased, beyond the value obtained for the sublethal dose with a very large cell-to-cell variation observed at *t*_1h_ (Fig. 6C-E). As expected, cell growth and division were rapidly arrested after this acute UV-C dose as evidenced by the flat growth curve of DR^WT^, DR^HUmCh^ and DR^HUmEos^ cultures (Fig. 5A and and SI Fig. S12B) and the stable cell size during stage 1*^L^* (SI Fig. S12C).

**Figure 5:**
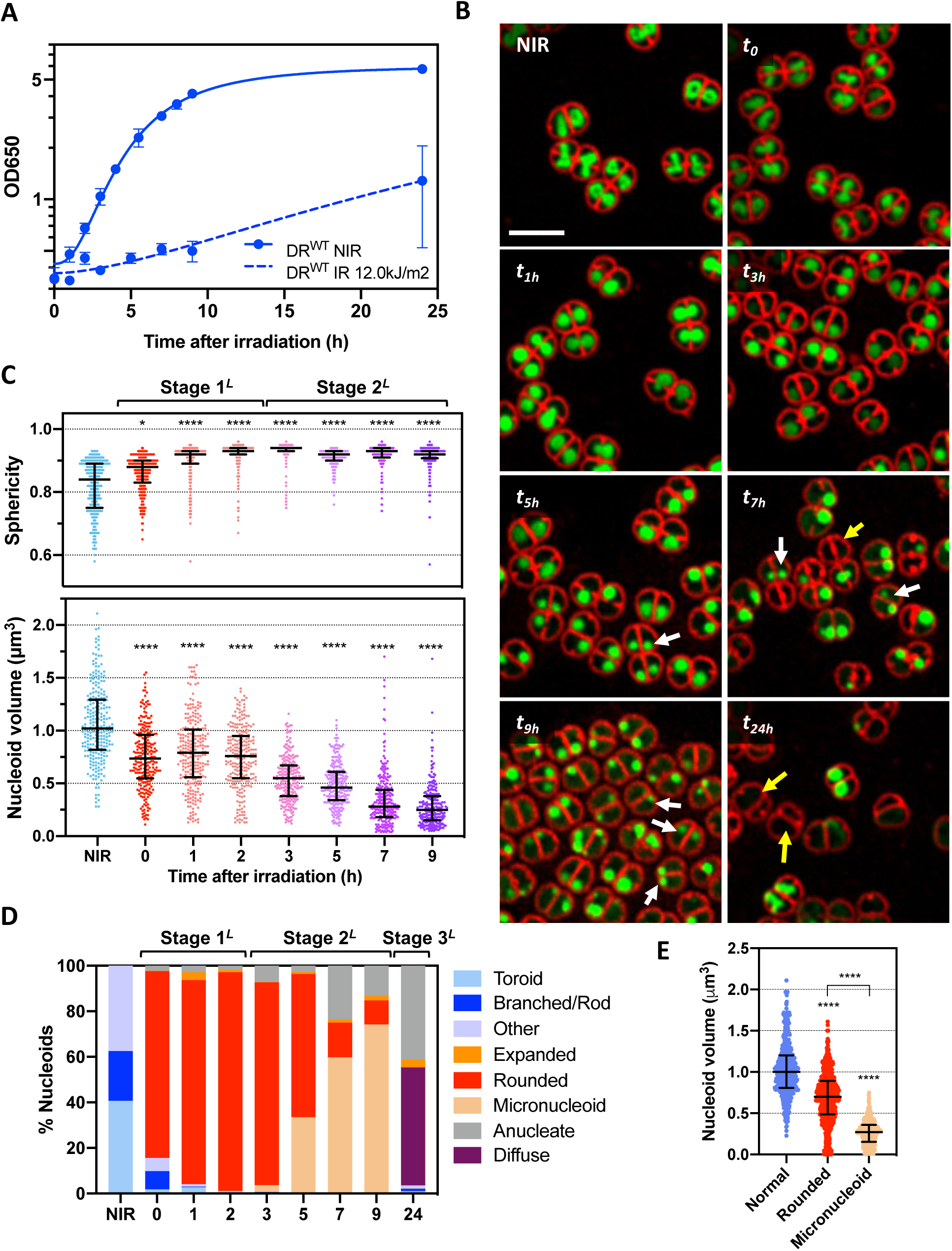
Effects of lethal UV-C exposure on nucleoid organization and morphology. (A) Growth curves of non-irradiated (NIR; full line) and irradiated DR^WT^ (12.0 kJ/m^2^; dashed line). Data represent the mean and standard deviation of at least 3 independent experiments. (B) Representative images of DR^WT^ cells stained with Syto9 and Nile Red at different timepoints after exposure to 12.0 kJ/m^2^ UV-C light. White and yellow arrows indicate cells with micro-nucleoids and anucleate cells respectively. Scale bar: 5 µm. (C) Nucleoid volume and sphericity of DR^WT^ cells at different timepoints after exposure to 12.0 kJ/m^2^ UV-C light (n=250). Error bars represent the median and interquartile range. * p<0.05, **** p<0.0001, Kruskal-Wallis statistical test performed in GraphPad Prism 8. (D) Evolution of nucleoid morphologies during the recovery from exposure to 12.0 kJ/m^2^ UV-C light (n>450). (C)-(D) The three different stages of the response phase are indicated above the plots. (E) Nucleoid volumes associated with different nucleoid morphologies (n>400). **** p<0.0001, Kruskal-Wallis statistical test performed in GraphPad Prism 8.

**Figure 6:**
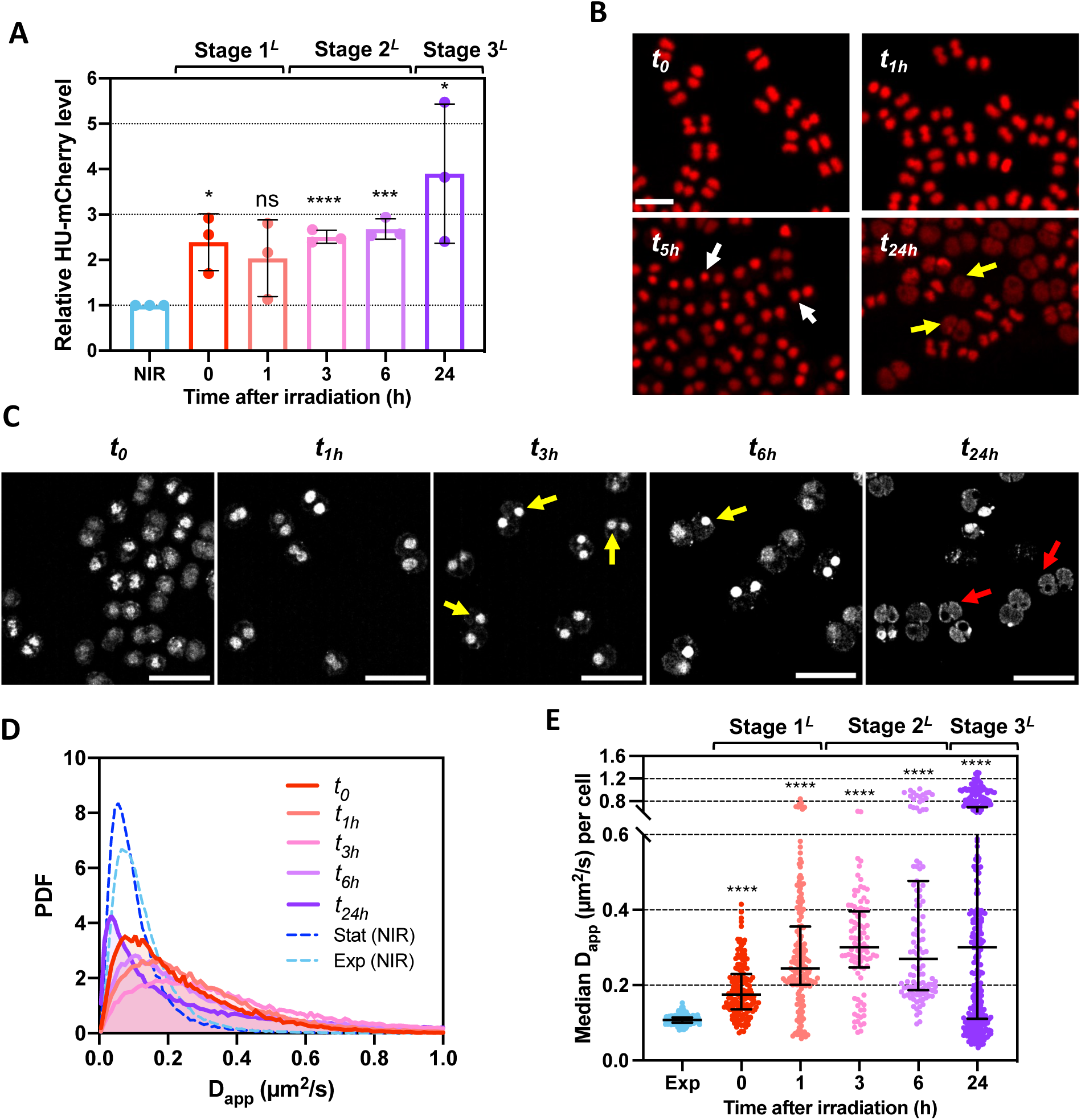
Effects of lethal UV-C exposure on the dynamics and abundance of HU. (A) Relative HU-mCherry levels determined by Western blot analysis at different timepoints after exposure to 12.0 kJ/m^2^ UV-C light (red to purple; n=3). The intensity of NIR samples was set to 1. Error bars represent the standard deviation. ns: non-significant, * p<0.05, *** p<0.001, **** p<0.0001, unpaired t-test performed in GraphPad Prism 8. (B) Representative images of DR^HUmCh^ at different timepoints after exposure to 12.0 kJ/m^2^ UV-C light. Micronucleoids are indicated with white arrows, while yellow arrows indicate cells with diffuse HU-mCherry staining. Scale bar: 5 µm. (C) Representative images of DR^HUmEos^ nucleoids at different timepoints after exposure to 12.0 kJ/m^2^ UV-C light. Yellow and red arrows indicate micronucleoids and diffuse nucleoids respectively. Scale bar: 5 µm. (D) Histograms of the apparent diffusion coefficient (D_app_) of HU-mEos4b in non-irradiated exponential DR^HUmEos^ (dashed blue line) cells and at different timepoints after exposure to 12.0 kJ/m^2^ UV-C light (red to purple; n>37). (E) Median D of HU-mEos4b per cell in non-irradiated exponential (light blue) DR^HU-mEos^ cells and at different timepoints after exposure to 12.0 kJ/m^2^ UV-C light (red to purple; n>37). Error bars represent the median and interquartile range. **** p<0.0001, Kruskal-Wallis statistical test performed in GraphPad Prism 8. (A)-(E) The three different stages of the response phase are indicated above the plots.

Whereas nucleoid compaction was reversed in stage 2*^SL^* after the sublethal dose, no such reversal was seen after the lethal dose. Instead, during stage 2*^L^* (*t*_2h_ to *t*_9h_ post-irradiation; Table S1), evidence for severe cellular dysfunction was observed: (i) lack of cell growth (SI Fig. S12C), (ii) increased fraction of cells exhibiting off-center nucleoids (Fig. 5B and SI Fig. S13B), (iii) accumulation of anucleate cells with reduced size due to a damaged cell wall (up to ∼25% of cells at *t_7h_*; Fig. 5B and D), (iv) stable HU levels exhibiting high HU diffusion (Fig. 6A and D), and (v) progressive reduction in the size of rounded nucleoids leading to the accumulation of micro-nucleoids with a mean volume of 0.27 µm^3^ corresponding to a 75% reduction in the nucleoid volume compared to non-irradiated samples (Fig. 5E).

In stage 3*^L^* (*t*_9h_ to *t*_24h_ post-irradiation; Table S1), there is a marked accumulation of cells exhibiting either no or diffuse staining of the DNA, suggesting that for a vast majority of cells, the integrity of the nucleoid was severely affected and genomic DNA was being heavily degraded (Fig. 5B-D and SI Fig. S13B). Indeed, at *t*_24h_, 95% of cells showed either no Syto9-DNA staining (40% of anucleate cells) or diffuse Syto9-DNA staining throughout the cytosol (55%). In accordance with this, HU diffusion remained at a high level throughout stages 2*^L^* and 3*^L^*, with many cells showing D_app_ values close or above 1 µm^2^/s, confirming the probable release of HU from the degraded genomic DNA in these cells (Fig. 6D-E). Strikingly, there were also many cells in which HU diffusion was very low (D_app_ < 0.1 µm^2^/s), possibly as a result of protein aggregation. Extensive DNA degradation was also evidenced in Syto9-stained DR^HUmCh^ cells, where an increasing number of HU-mCherry positive and Syto9 negative nucleoids were seen to accumulate during stage 3*^L^* (SI Fig. S14).

## DISCUSSION

In this study, we have investigated nucleoid remodeling in *D. radiodurans* in response to two different types of stress: (i) entry into stationary phase, which is associated with nutritional stress and (ii) exposure to UV-C light, which is known to cause severe DNA damage. A comparison of the response to these two types of stress (Fig. 7) provides important insight into the mechanisms underlying bacterial nucleoid remodeling.

**Figure 7:**
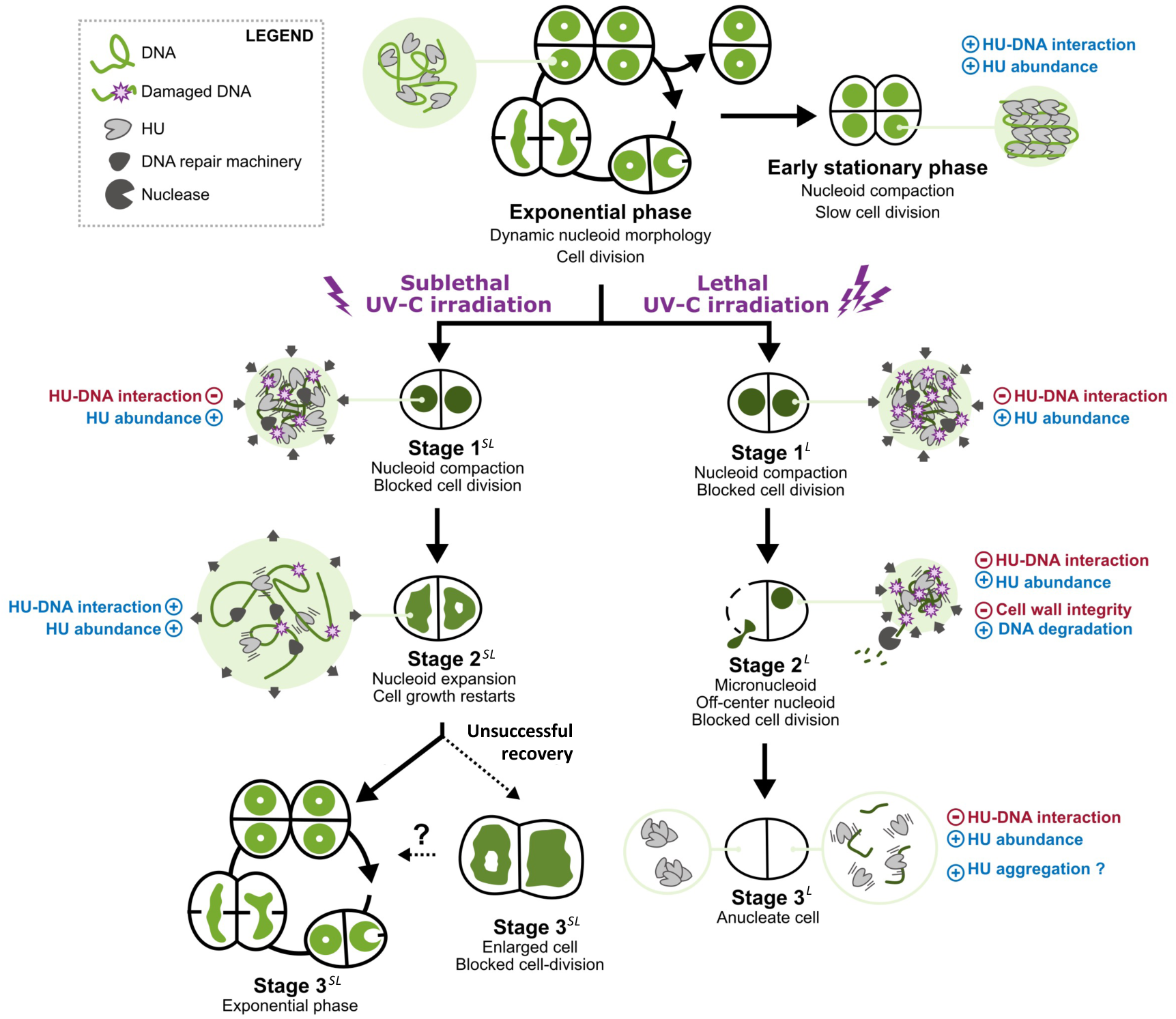
Model of stress-induced nucleoid remodeling in. D. radiodurans. Schematic diagram summarizing the main stages of nucleoid remodeling in *D. radiodurans* induced by either entry into stationary phase (top right) or by exposure to either sublethal (lower left) or lethal (lower right) doses of UV-C irradiation. Changes in nucleoid size and shape are accompanied by significant variations in the level and dynamics of the HU protein, likely reflecting altered HU-DNA interactions. Nucleoid remodeling is also tightly coordinated with cell growth and division.

In both cases, nucleoids rapidly became more compact and changed their morphology to adopt a more rounded shape, and this phenomenon was associated with increased levels of the HU protein, in agreement with earlier work that showed that conversely HU depletion leads to nucleoid expansion (Nguyen *et al*., 2009). Upon entry into stationary phase, however, this nucleoid compaction was accompanied by a significant decrease in HU diffusion (Fig. 7), suggesting that HU is binding more tightly, and perhaps in a different configuration, to the genomic DNA compared to exponentially growing cells. The increased HU levels in stationary phase cells combined with the markedly reduced volumes of stationary phase nucleoids is likely to result in a strongly increased local concentration of HU in such cells. This may indeed modify the DNA binding properties of HU and favor the formation of HU-DNA assemblies in which HU polymerizes along the genomic DNA, thereby stiffening and bundling the genomic DNA (Fig. 7). A similar model was recently proposed by Hammel and colleagues for *E. coli* HU in stationary phase (Hammel *et al*., 2016; Remesh *et al*., 2020). This model is also supported by *in vitro* atomic force microscopy studies of DrHU-plasmid DNA assemblies, in which stiffening and bridging of plasmid DNA by DrHU were observed at high HU:DNA ratios only (Chen *et al*., 2020) and would explain the reduced apparent diffusion of HU observed in the highly condensed stationary phase *D. radiodurans* nucleoids. By maintaining this condensed genomic configuration in stationary phase, HU also likely serves as a global transcriptional regulator, facilitating the transcription of stationary phase specific genes required for survival under resource-limited growth conditions as has been reported in *E. coli* (Hengge, 2011; Sobetzko *et al*., 2012; Lal *et al*., 2016).

As in stationary phase cells, exposure of *D. radiodurans* cells to UV-C light rapidly induced nucleoid compaction, although to a lesser extent, and this phenomenon was again associated with an increase in HU levels regardless of the UV dose. HU thus appears to be a key player in the process of stress-induced nucleoid condensation and regulated *de novo* synthesis of HU during the very early stage of nucleoid remodeling may contribute to maintaining the integrity of the nucleoid by providing functional HU protein (Fig. 7). However, while HU diffusion was decreased in stationary phase cells, it was significantly increased in response to UV-C light. The structure of condensed stationary phase nucleoids is thus distinct from that of UV-C-irradiated nucleoids, although in both cases, HU-DNA interactions are likely playing an important role in the underlying organization. Increased mobility of the irradiated genomic DNA due to DNA damage could have been one of the causes of the observed increased HU dynamics, but the tracking of the *oriC* and *ter* loci of the major chromosome of *D. radiodurans* showed no evidence for a substantial change in DNA mobility after exposure to UV-C light. The increased HU diffusion thus most likely results from weakened HU-DNA interactions. These may be induced by either post-translational modifications and/or physico-chemical changes occurring at the level of HU and/or the genomic DNA, directly induced by the high-intensity UV-C light and the overproduction of ROS (Kehrer, 2000; Díaz-Riaño *et al*., 2019).

A major consequence of HU release from the genomic DNA as a result of weakened HU-DNA interactions could be compaction or collapse of the nucleoid (Fig. 7), although additional factors are likely also involved in this process. UV-induced arrest (or partial arrest) of cellular processes, including cell growth and division, but also DNA replication and transcription or protein translation, which are known to induce changes in nucleoid organization (Woldringh *et al*., 1995; Miangolarra *et al*., 2021), is likely to also contribute to this rapid stress-induced nucleoid condensation. Similarly, reduced transcription and translation as well as severe damage to the bacterial cytoskeleton network may be responsible for the subsequent drift of nucleoids off-center, that was particularly striking in cells exposed to lethal UV-C light (Fig. 7). Using theoretical approaches, Marc Joyeux showed that bacterial nucleoids preferentially locate to the edges of the cell, where the curvature of the cell wall is the highest (Joyeux, 2019), suggesting that in healthy cells active cellular processes, including transcription and translation (Miangolarra *et al*., 2021), are indeed responsible for maintaining the nucleoid in a central position. These processes are likely impaired in the strongly irradiated cells, causing the nucleoid to lose its structure and drift to the edge of the cell.

A major difference in the response of *D. radiodurans* to sublethal versus lethal UV-C doses was the reversibility of the process. In *D. radiodurans* cells exposed to sublethal UV-C light, the rapid initial nucleoid compaction stage (stage 1*^SL^*) was followed by a slower decompaction phase (stage 2*^SL^*) during which the nucleoids progressively recovered their well-structured states and cells recovered their ability to grow and divide. HU is likely to play an important role in this recovery stage (Fig. 7). We, indeed, observed that the kinetics of recovery of nucleoid morphology after exposure to sublethal UV-C light coincided with that of recovery of HU dynamics, suggesting that progressive re-assembly of HU on the genomic DNA is an important determinant of nucleoid organization and may be critical for regulating access of DNA repair enzymes to the genomic DNA. On the contrary, in stage 2*^L^*, a lethal dose of UV-C light lead to progressive loss of nucleoid structure, heavy degradation of the genomic DNA, full arrest of cell growth and division, and eventually to the accumulation of damaged and anucleate cells in stage 3*^L^*. In such cells, HU diffusion was found to be either very high, indicating that HU-DNA interactions were largely disrupted, or instead very low, possibly as a result of HU aggregation in the absence of genomic DNA on which to assemble.

Cell survival is thus essential to restore nucleoid organization following exposure to UV-C light and in contrast to nucleoid condensation which may be a mostly passive process, nucleoid decompaction, and more generally cell recovery, largely relies on active cellular processes (Fig. 7). In particular, active checkpoint control factors must be functional to rapidly coordinate the arrest of the cell cycle and chromosome segregation with the intervention of the DNA repair machinery to restore the integrity of the genome. A similar sequential compaction and expansion of nucleoids was reported in UV-C irradiated *E. coli* cells before cell division could restart (Odsbu and Skarstad, 2014; Estévez Castro *et al*., 2018). Obsbu and Skarstad proposed that the initial phase of compaction serves to stabilize the DNA and repair DNA double-strand breaks, while repair of other lesions takes place largely during the decompaction stage (Odsbu and Skarstad, 2014). In *D. radiodurans*, nucleoid condensation appears to occur regardless of the nature of the stress and its intensity, suggesting that it may constitute an early stress response mechanism, which could act mostly as a protective measure to minimize dispersion of the damaged genome. Earlier studies have shown that activation of the DNA repair machinery occurs 30 to 60 minutes after exposure to radiation (Liu *et al*., 2003; Basu and Apte, 2012), suggesting that most of the repair process would therefore take place during the decompaction stage. This hypothesis is in line with studies in eukaryotic cells showing that the chromatin decondenses in response to DNA damage to facilitate repair (Hauer *et al*., 2017; dos Santos *et al*., 2020). The 2-3h delay in the growth of *D. radiodurans* after exposure to UV-C is also in good agreement with this hypothesis and the estimated time needed to allow cells to repair heavy radiation-induced DNA damage (Zahradka *et al*., 2006).

In *D. radiodurans,* although many cells recovered well from exposure to the sublethal UV-C dose and started to grow and divide again during stage 2*^SL^* of the recovery phase, a fraction of cells exhibited highly expanded nucleoids. These cells were typically larger in size and lacking a division septum, suggesting that they had recovered their ability to grow, but not to divide. Castro *et al*. showed that *E. coli* cells that are deficient in DNA repair proteins reach the expanded nucleoid stage, but are unable to restart cell division after UV-C irradiation, suggesting that DNA repair is required before cell division can restart (Estévez Castro *et al*., 2018). *D. radiodurans* cells exhibiting highly expanded nucleoids were thus probably either failing to repair their DNA damage or delayed in their repair process, leading to arrested cell division and partial loss of their nucleoid organization. A fraction of such cells may eventually recover and start dividing again after completion of the repair process. Interestingly, such cells arose progressively during the recovery, suggesting that cells that initially appeared to have recovered subsequently developed cellular defects probably as a consequence of accumulated DNA damage.

In conclusion, this study highlights the key role of HU and the versatility of its interactions with genomic DNA in the nucleoid remodeling process in *D. radiodurans* and the survival strategy of this remarkable organism. However, besides HU, additional factors, such as DdrC, a *Deinococcus*-specific NAP that is strongly up-regulated in response to irradiation (Bouthier de la Tour *et al*., 2017; Banneville *et al*., 2022), the partitioning factors (ParA/ParB) or the DNA gyrase, all of which are found in nucleoids after irradiation (Bouthier de la Tour *et al*., 2013), may also contribute to varying extent to this nucleoid remodeling process. Further work will be needed to explore these different avenues. Moreover, a number of recent studies have revealed the importance of post-translational modifications of NAPs for regulating the activities and DNA-binding properties of these small basic proteins, so in the future, establishing the post-translational modification profiles of key NAPs, and notably of HU, at different stages of the nucleoid remodeling process will certainly shed new light on the underlying mechanisms.

## MATERIALS AND METHODS

### Bacterial cultures

Bacterial strains used in this study are listed in Table S2. All *D. radiodurans* strains were derivatives of the wild-type strain R1 ATCC 13939 (DR^WT^). The genetically engineered strain of *D. radiodurans* expressing HU fused to mEos4b (GY16993; DR^HUmEos^) was obtained by the tripartite ligation method (Mennecier *et al*., 2004). For this, a codon-optimized gene encoding mEos4b to which we added the 9 N-terminal residues of mCherry was synthesized and subloned into plasmid pEX-128 by Eurofins MWG Biotech (SI Fig. S15). Obtaining a viable and functional strain of *D. radiodurans* expressing HU fused to a fluorescent protein is not trivial. Since we knew that mCherry fusions were functional, we decided to add the extra N-terminal residues of mCherry to the N-terminus of mEos4b. We inserted a hygromycin resistance (*hph*) cassette into the AgeI/XbaI sites of pEX-128 downstream of the mEos4b gene, then amplified the mEos4bΩ*hph* cassette and the regions flanking the insertion site (3’ end of *hu* gene, DR_A0065) by PCR using oligonucleotides listed in Table S3, and after restriction digestion the three fragments were ligated together. *D. radiodurans* cells were then transformed by the ligation product and plated on selective medium containing 50 µg/ml hygromycin, leading to allelic replacement on one genome copy. Because *D. radiodurans* is multigenomic, the transformant colonies were further streaked three times successively on selective medium to ensure that all copies of the genome had incorporated the foreign DNA. This was then confirmed by PCR analysis and DNA sequencing, which also revealed a point mutation in mEos4b (Ala^112^ to Val^112^). *D. radiodurans* cells were grown aerobically at 30°C in a shaking incubator (160 rpm) in TGY2X medium (Bonacossa *et al*., 2002). The medium was supplemented with the appropriate antibiotics: 3.4 μg/ml chloramphenicol for DR^HUmCh^ GY15743 strain and *oriC*/*ter* labelled strains GY15787 and GY15800, and 50 µg/ml hygromycin for DR^HUmEos^ GY16993. Typically for microscopy experiments, *D. radiodurans* cells were pre-grown the day before and then diluted for an overnight growth until reaching exponential (OD_650_ ∼0.3-0.5) or early stationary phase (OD_650_ >3) the next morning. *oriC*/*ter* labelled strains GY15787 and GY15800 were grown overnight and were diluted 60X the next morning in TGY2X and grown for a further 5h until reaching an OD_650_ between 0.3 and 0.5. *Escherichia coli* strain DH5α was grown at 37°C in LB medium. Optical density measurements were made on a Clariostar (BMG Labtech) plate reader.

### UV irradiation and survival curve

Before irradiation, cells were washed two times with minimal medium M9DR (45.5 mM Na_2_HPO_4_, 22 mM KH_2_PO_4_ and 15.1 mM (NH_4_)_2_SO_4_, pH7.3) to eliminate the strongly UV-absorbing TGY2X medium. Cells were finally resuspended in M9DR medium at an OD_650_ between 0.3 and 0.5 and transferred to a 6-well plate (1 ml per well). Cells were irradiated at a low cell density and in a minimal volume to ensure a homogeneous exposure of the cells. UV-C (254nm) irradiation was performed using a Stratalinker oven equipped with germicide UV-C lamps producing a continuous dose of ∼30J/m^2^/sec. Cells were typically irradiated between 0.5 and 5 minutes. The exact dose received by the cells was determined using a UV-C light monitor placed next to the 6-well plate in the UV oven. Exposure to UV-C irradiation did not affect the fluorescence of the fluorescent proteins (SI Fig. S3B). After irradiation, cells were centrifuged and resuspended in fresh TGY2X medium. For the recovery experiments, these cultures were returned to the 30°C shaking incubator. Samples were collected immediately after resuspending cells in TGY2X medium (*t*_0_) and then 1, 2, 3, 5, 7, 9 and 24 hours post-irradiation (for the confocal analysis) or 1, 3, 6 and 24 hours post-irradiation (for the spt analysis). OD_650_ measurements were made every hour on three independent cultures for 9h post-irradiation and then the following day (*t*_24h_). Full UV-C survival curves were performed for *E. coli* DH5α and exponential and stationary phase DR^WT^ by serially diluting non-irradiated and irradiated cultures (exposed to 0 to 12 kJ/m^2^ for DR^WT^ or 0 to 0.95 kJ/m^2^ for *E. coli*) and spotting 8 µl of each dilution on TGY1X or LB (for *E. coli*) agar plates. After 48h incubation of the plates at 37°C, individual colonies were counted and the surviving fraction was determined using non-irradiated cells as a reference. A similar procedure was used to determine the survival fraction of DR^HU-mCh^ and DR^HU-mEos^ strains after exposure to 1.9 and 12.0 kJ/m^2^.

### Sample preparation for confocal microscopy and sptPALM

For confocal microscopy analysis, DR^WT^ cells were stained with Nile Red (30 µM) and Syto9 (200 nM) for 10 min at room temperature, to visualize the cell membrane and nucleoid, respectively, while DR^HU-mCh^ cells were only stained with syto9 (200 nM). The cells were then harvested by centrifugation and resuspended in 10 μl TGY2X. 0.5 μl of this cell suspension was then placed on a 1.5% (w/v) low melting agarose (LMA; Bio-Rad) TGY2X pad prepared on a cover glass using a gene frame (Thermo Scientific). A 2 mm-wide band of LMA was cut out to provide oxygen. A glass slide was then placed on top of the pad and the sample was sealed with picodent twinsil. Sample preparation for microscopy typically took approximately 30 min, so cells were usually imaged 30 min after the designated timepoint. The samples were maintained at 30°C during image acquisition. For spt analysis, cells were washed 3 times in M9DR medium to limit autoblinking, as described previously (Floc’h *et al*., 2018), and were finally resuspended in 20-100 µl M9DR medium, depending on the size of the pellet. 10 µl of this cell suspension was then placed on a coverslip cleaned with an ozone cleaner device (UVOCS) and the cells were left to sediment for 2 minutes, after which the excess liquid was removed. After a further 2-4 minute of air-drying, 10 µl 1.5% (w/v) LMA in M9DR was poured over the cells and a glass slide was placed on top to evenly distribute the LMA. Samples were sealed with picodent Twinsil® and imaged within 20 minutes.

### Confocal data acquisition

Spinning-disk confocal microscopy was performed using an Olympus IX81 inverted microscope equipped with a Yokogawa CSU-X1 confocal head. The excitation laser beam (Ilas2 laser bench, GATACA systems) was focused to the back focal plane of a 100X 1.49-numerical-aperture (NA) oil immersion apochromatic objective. For cell and nucleoid analyses, series of Z-planes were acquired every 100 nm using a PRIOR N400 piezo stage. Fluorescence excitation was performed at 488 nm for Syto 9 and 561 nm for Nile Red or mCherry. Fluorescence emission was collected with an Andor iXon Ultra EMCCD camera through a quad-band Semrock™ Di01-T405/488/568/647 dichroic mirror and single-band emission filters adapted to each fluorophore used: 520 nm for Syto9 (FF02-520/28 Semrock™), and 600 nm for mCherry or Nile Red (ET600/50m Chroma™). Data acquisition was performed using Metamorph 7.10 (Molecular devices).

### SptPALM data acquisition and tracking of oriC/ter loci

SptPALM acquisitions on the DR^HUmEos^ strain were acquired on a SAFE 360 (Abbelight) SMLM set up. Data was acquired at 27°C under continuous illumination with 400 W/cm² 561 nm light and a frame time of 10 ms. The intensity of the 405 nm laser was manually increased during acquisition to maintain approximately constant localization density. 40,000 – 60,000 frames were acquired per dataset. A transmission light image was acquired prior to sptPALM acquisition for segmentation of the cells. *oriC* and *ter* labeled strains of *D. radiodurans* were imaged on a home-built PALM set-up. First, a bright field image and an image of HU-mCherry were collected (100 ms exposure time, 10 W/cm^2^ 561 nm light). Second, the *oriC* or *ter* sites labelled with ParB-GFP were imaged every second using an exposure time of 100 ms and 10 W/cm^2^ 488 nm light, until all sites had bleached (∼100-500 frames).

### Confocal data analysis

3D nucleoids were segmented by automatic thresholding of the Syto9 or mCherry fluorescence signal using the ‘surface’ procedure of the Imaris software (Oxford instruments) after parameter optimization (surface smoothing detail = 0.1 µm and background subtraction = 0.2 µm). The volume, sphericity and mean channel intensities (in the two channels) of the segmented nucleoids were then extracted for further analysis. After exposure to sublethal and lethal doses of UV-C light, 450-550 individual cells from each timepoint (*t*_0_ to *t*_24h_) were classified in terms of phase of the cell cycle (phase P1 to P6 for exponentially growing cells and phases P1 to P8 for stationary cells), cell perimeter, cell integrity, nucleoid volume, nucleoid morphology and nucleoid position (centered or off-center). Cell perimeters were obtained by automatic segmentation of cells using Cellpose (Stringer *et al*., 2021) and conversion of the obtained masks into ROIs in Fiji (Schneider *et al*., 2012), which could be used to extract the perimeter of the segmented cells. Phase 5 cells, in particular, but also some distorted cells were poorly segmented by Cellpose. In these cases, segmentation was performed manually in ImageJ.

### sptPALM and oriC/ter tracking data analysis

Nucleoids were manually segmented from an overlay of the bright field and super-resolved images of HU-mEos4b. A small fraction of cells with poorly defined nucleoids were excluded from further analysis. Despite the washing steps, occasionally a fraction of cells still showed strong autoblinking (Floc’h *et al*., 2018); these cells were also discarded from the analysis. Single molecules were localized using the Thunderstorm plugin in ImageJ (Ovesný *et al*., 2014). Trajectories of single molecules were obtained using the tracking software Swift (Endesfelder *et al*. in prep), using the following tracking parameters: ‘exp_displacement = 60 nm’, ‘p_bleach = 0.1‘, ‘p_blink = 0.2’, ‘p_reappear = 0.5’, ‘precision = 22 nm’, ‘max_displacement = 250 nm’, ‘max_displacement_pp = 250 nm’ and ‘max_blinking_duration = 2 frames’. Default values were used for the other parameters. Swift splits tracks, which contain motion changes (i.e. immobile -> diffusive), into segments containing unique behaviors. These segments were considered as trajectories for further analysis. Of note, only a small fraction of the tracks of HU-mEos4b contained multiple segments (∼1-2%) suggesting that the time HU-mEos4b spends in a given state is relatively long compared to the average track length.

Trajectories were further analyzed in MATLAB to assign them to specific nucleoid morphologies based on the applied segmentation and to calculate the apparent diffusion coefficients. Apparent diffusion coefficients of single trajectories (Di*) were calculated from the 1-step mean squared jump distance (MJD) as described by Stracy et al. (Stracy *et al*., 2021) using:

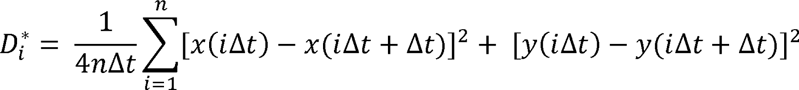

where *n* is the number of displacements over which the 1-step MSD is calculated, *x*(*t*) and *y*(*t*) are the coordinates of the molecule at time t and Δ*t* is the frametime. In this work, 4 displacements per track (made of at least 5 measurements) were taken into account (n = 4).

*OriC* and *ter* loci were localized using Thunderstorm (Ovesný *et al*., 2014). Trajectories were obtained using Swift (Endesfelder *et al*. in prep) with the following tracking parameters: ‘exp_displacement = 50 nm’, ‘p_bleach = 0.001 ‘, ‘precision = 50 nm’, ‘max_displacement = 250 nm’, ‘max_displacement_pp = 250 nm’. Default values were used for the other parameters. Trajectories were further analyzed in MATLAB to calculate the apparent diffusion coefficients. Only trajectories with >10 localizations were considered.

### Western blot analysis

10 ml samples of DR^HUmCh^ cultures were collected immediately after resuspending cells in TGY2X medium (t_0_) and then 1, 2 (for sublethal dose only), 3, 6 and 24 hours post-irradiation with either sublethal (1.9 kJ/m^2^) or lethal (12.0 kJ/m^2^) UV-C dose. Similarly, non-irradiated samples of exponentially and stationary phase cells were also processed alongside these irradiated samples. Each culture was resuspended in 0.2 ml PBS supplemented with protease inhibitors (Roche), DNaseI (Roche) and lysozyme (Roche). Cell suspensions were lysed by two 10-sec sonications with a microtip followed by 45 min mechanical shaking with 100 µl glass beads (Macherey Nagel) at 4°C. Cell debris were eliminated by centrifugation at 14,000 rpm for 10 min and the soluble fraction was recovered. 7 µg of each cell extract was loaded on a 12% stain-free SDS-PAGE gel (BioRad) and separated by gel electrophoresis at 175 V for 45 min, before transferring the bands to a nitrocellulose membrane using the BioRad transblot system. A stain-free image of the blot was acquired at this point for normalization of the lanes before blocking the membrane with 5% milk in PBS for 30 min. The membrane was then incubated overnight at 4°C with a mouse monoclonal anti-mCherry antibody (Origene; 1:1500) diluted in 5% milk in PBS. After three washes with PBS supplemented with 0.02% Tween20 (PBS-Tween), the membrane was further incubated 1 hour at 25°C with a second antibody (anti-mouse HRP conjugated antibody; Sigma) diluted 1:10,000 in PBS-Tween. After three additional washes with PBS-Tween, the bands were revealed by electrochemiluminescence (ECL) using Clarity Western substrate (BioRad) on a Chemidoc MP imager (BioRad). Using ImageLab (BioRad), bands corresponding to HU-mCherry were quantified relative to the non-irradiated sample loaded on the same gel after normalization of the lanes using the stain-free image. Blots were performed in triplicate and the mean and standard deviation of the obtained values were plotted in GraphPad Prism 8. Statistical differences were determined using a one-way ANOVA (Holm Sidak) multiple comparisons test.

### Statistical analysis

Unless otherwise stated, non-parametric Kruskal-Wallis statistical tests were performed with GraphPad Prism 8 in order to assess the significance of the data. Two-tailed p values below 0.05 were considered as significant and were indicated with asterisks: ns: non-significant, * p ≤0.05, ** p ≤0.01, *** p ≤0.001, **** p ≤0.0001.

## ACKNOWLEDGEMENTS

We thank Claire Bouthier de la Tour for help in the preparation of the genetically engineered strain of *D. radiodurans*. This work was supported by the CEA Radiobiology program. JW’s PhD position was funded by GRAL, a project of the University Grenoble Alpes graduate school (Ecoles Universitaires de Recherche) CBH-EUR-GS (ANR-17-EURE-0003). This work was supported by the Agence Nationale de la Recherche (grants no. ANR-20-CE11-0013-01 and ANR-22-CE11-0029-01) and used the M4D imaging platform of the Grenoble Instruct-ERIC Center (ISBG : UMS 3518 CNRS-CEA-UGA-EMBL) within the Grenoble Partnership for Structural Biology (PSB), supported by FRISBI (ANR-10-INBS-05-02) and GRAL (ANR-10-LABX-49-01), financed within the University Grenoble Alpes graduate school (Ecoles Universitaires de Recherche) CBH-EUR-GS (ANR-17-EURE-0003). IBS acknowledges integration into the Interdisciplinary Research Institute of Grenoble (IRIG, CEA).

## AUTHOR CONTRIBUTIONS

PV, JW, JPK, DB and JT designed the research. PV, JW, FL and JPK performed the microscopy experiments. PS prepared the genetically modified strains of *D. radiodurans*. PV, JW, JPK, FC, DB and JT analyzed the data. PV, JW, DB and JT wrote the manuscript and all authors discussed the results and approved the manuscript.

## REFERENCES

Aki, T. (1997) Repressor induced site-specific binding of HU for transcriptional regulation. The EMBO Journal 16: 3666–3674.

Azam, T.A., and Ishihama, A. (1999) Twelve Species of the Nucleoid-associated Protein from *Escherichia coli*. Journal of Biological Chemistry 274: 33105–33113.

Banneville, A.-S., Bouthier de la Tour, C., De Bonis, S., Hognon, C., Colletier, J.-P., Teulon, J.-M., et al. (2022) Structural and functional characterization of DdrC, a novel DNA damage-induced nucleoid associated protein involved in DNA compaction. Nucleic Acids Research 50: 7680–7696 10.1093/nar/gkac563. Accessed August 16, 2022.

Basu, B., and Apte, S.K. (2012) Gamma radiation-induced proteome of *Deinococcus radiodurans* primarily targets DNA repair and oxidative stress alleviation. Mol Cell Proteomics 11: M111.011734.

Battista, J.R. (1997) Against all odds: the survival strategies of *Deinococcus radiodurans*. Annu Rev Microbiol 51: 203–24.

Bettridge, K., Verma, S., Weng, X., Adhya, S., and Xiao, J. (2021) Single-molecule tracking reveals that the nucleoid-associated protein HU plays a dual role in maintaining proper nucleoid volume through differential interactions with chromosomal DNA. Molecular Microbiology 115: 12–27 10.1111/mmi.14572. Accessed June 20, 2023.

Boor, K.J. (2006) Bacterial Stress Responses: What Doesn’t Kill Them Can Make Them Stronger. PLoS Biol 4: e23.

Bouthier De La Tour, C., Armengaud, J., Dulermo, R., Blanchard, L., Devigne, A., De Groot, A., et al. (2015) The abundant and essential HU proteins in *Deinococcus deserti*and *Deinococcus radiodurans* are translated from leaderless mRNA. Microbiology 161: 2410–2422.

Bouthier de la Tour, C., Mathieu, M., Meyer, L., Dupaigne, P., Passot, F., Servant, P., et al. (2017) In vivo and in vitro characterization of DdrC, a DNA damage response protein in *Deinococcus radiodurans* bacterium. PLoS One 12: e0177751.

Bouthier de la Tour, C., Passot, F.M., Toueille, M., Mirabella, B., Guerin, P., Blanchard, L., et al. (2013) Comparative proteomics reveals key proteins recruited at the nucleoid of *Deinococcus* after irradiation-induced DNA damage. Proteomics 13: 3457–69.

Boutte, C.C., and Crosson, S. (2013) Bacterial lifestyle shapes stringent response activation. Trends in Microbiology 21: 174–180.

Chawla, M., Mishra, S., Anand, K., Parikh, P., Mehta, M., Vij, M., et al. (2018) Redox-dependent condensation of the mycobacterial nucleoid by WhiB4. Redox Biology 19: 116–133.

Chen, S.W.W., Banneville, A.S., Teulon, J.M., Timmins, J., and Pellequer, J.L. (2020) Nanoscale surface structures of DNA bound to *Deinococcus radiodurans* HU unveiled by atomic force microscopy. Nanoscale 12: 22628–22638.

Cox, M.M., and Battista, J.R. (2005) *Deinococcus radiodurans*-The consummate survivor. Nature Reviews Microbiology 3: 882–892.

Cuypers, M.G., Mitchell, E.P., Romao, C.V., and McSweeney, S.M. (2007) The crystal structure of the Dps2 from *Deinococcus radiodurans* reveals an unusual pore profile with a non-specific metal binding site. J Mol Biol 371: 787–99.

Dame, R.T., Rashid, F.-Z.M., and Grainger, D.C. (2020) Chromosome organization in bacteria: mechanistic insights into genome structure and function. Nature Reviews Genetics 21: 227–242 10.1038/s41576-019-0185-4.

De Zitter, E., Thédié, D., Mönkemöller, V., Hugelier, S., Beaudouin, J., Adam, V., et al. (2019) Mechanistic investigation of mEos4b reveals a strategy to reduce track interruptions in sptPALM. Nature Methods 16: 707–710 10.1038/s41592-019-0462-3.

Díaz-Riaño, J., Posada, L., Acosta, I.C., Ruíz-Pérez, C., García-Castillo, C., Reyes, A., and Zambrano, M.M. (2019) Computational search for UV radiation resistance strategies in *Deinococcus swuensis* isolated from Paramo ecosystems. PLoS ONE 14.

Dillon, S.C., and Dorman, C.J. (2010) Bacterial nucleoid-associated proteins, nucleoid structure and gene expression. Nat Rev Microbiol 8: 185–195.

Dilweg, I.W., and Dame, R.T. (2018) Post-translational modification of nucleoid-associated proteins: an extra layer of functional modulation in bacteria? Biochemical Society Transactions 46: 1381–1392 10.1042/BST20180488. Accessed March 3, 2022.

dos Santos, Á., Cook, A.W., Gough, R.E., Schilling, M., Olszok, N.A., Brown, I., et al. (2020) DNA damage alters nuclear mechanics through chromatin reorganization. Nucleic Acids Res 49: 340–353.

Dworkin, J., and Harwood, C.S. (2022) Metabolic Reprogramming and Longevity in Quiescence. Annu Rev Microbiol 76: 91–111.

Endesfelder, M., Schließl, C., Turkowyd, B., Lechner, T., and Endesfelder, U. swift – fast, probabilistic tracking for dense, highly dynamic single-molecule data. Manuscript in prep.

Estévez Castro, C.F., Serment-Guerrero, J.H., and Fuentes, J.L. (2018) Influence of uvrA, recJ and recN gene mutations on nucleoid reorganization in UV-treated *Escherichia coli* cells. FEMS Microbiology Letters 365: fny110 10.1093/femsle/fny110. Accessed August 16, 2023.

Floc’h, K., Lacroix, F., Barbieri, L., Servant, P., Galland, R., Butler, C., et al. (2018) Bacterial cell wall nanoimaging by autoblinking microscopy. Sci Rep 8: 14038 http://www.nature.com/articles/s41598-018-32335-z. Accessed October 19, 2019.

Floc’h, K., Lacroix, F., Servant, P., Wong, Y.-S., Kleman, J.-P., Bourgeois, D., and Timmins, J. (2019) Cell morphology and nucleoid dynamics in dividing *Deinococcus radiodurans*. Nature Communications 10: 3815 10.1038/s41467-019-11725-5.

Gefen, O., Fridman, O., Ronin, I., and Balaban, N.Q. (2014) Direct observation of single stationary-phase bacteria reveals a surprisingly long period of constant protein production activity. Proc Natl Acad Sci USA 111: 556–561.

Ghosh, S., and Grove, A. (2004) Histone-like protein HU from *Deinococcus radiodurans* binds preferentially to four-way DNA junctions. J Mol Biol 337: 561–71.

Ghosh, S., and Grove, A. (2006) The *Deinococcus radiodurans* -Encoded HU Protein Has Two DNA-Binding Domains. Biochemistry 45: 1723–1733.

Ghosh, S., Padmanabhan, B., Anand, C., and Nagaraja, V. (2016) Lysine acetylation of the *Mycobacterium tuberculosis* HU protein modulates its DNA binding and genome organization. Molecular Microbiology 100: 577–588.

Grove, A. (2011) Functional evolution of bacterial histone-like HU proteins. Curr Issues Mol Biol 13: 1–12.

Gupta, M., Sajid, A., Sharma, K., Ghosh, S., Arora, G., Singh, R., et al. (2014) HupB, a Nucleoid-Associated Protein of *Mycobacterium tuberculosis*, Is Modified by Serine/Threonine Protein Kinases *In Vivo*. J Bacteriol 196: 2646–2657.

Haikarainen, T., and Papageorgiou, A.C. (2010) Dps-like proteins: structural and functional insights into a versatile protein family. Cell Mol Life Sci 67: 341–351.

Hammel, M., Amlanjyoti, D., Reyes, F.E., Chen, J.H., Parpana, R., Tang, H.Y.H., et al. (2016) HU multimerization shift controls nucleoid compaction. Science Advances 2.

Harsojo, Kitayama, S., and Matsuyama, A. (1981) Genome multiplicity and radiation resistance in *Micrococcus radiodurans*. J Biochem 90: 877–80.

Hauer, M.H., Seeber, A., Singh, V., Thierry, R., Sack, R., Amitai, A., et al. (2017) Histone degradation in response to DNA damage enhances chromatin dynamics and recombination rates. Nat Struct Mol Bio l24: 99–107.

Hengge, R. (2011) Stationary-Phase Gene Regulation in *Escherichia coli*. EcoSal Plus 4: ecosalplus.5.6.3.

Hołówka, J., and Zakrzewska-Czerwińska, J. (2020) Nucleoid Associated Proteins: The Small Organizers That Help to Cope With Stress. Front Microbiol 11: 590.

Hou, J., Dai, J., Chen, Z., Wang, Y., Cao, J., Hu, J., et al. (2022) Phosphorylation Regulation of a Histone-like HU Protein from *Deinococcus radiodurans*. Protein & Peptide Letters 29: 891–899 http://www.eurekaselect.com/article/125634.

Huisman, O., Faelen, M., Girard, D., Jaffé, A., Toussaint, A., and Rouvière-Yaniv, J. (1989) Multiple defects in Escherichia coli mutants lacking HU protein. J Bacteriol 171: 3704–3712.

Joyeux, M. (2019) Preferential Localization of the Bacterial Nucleoid. Microorganisms 7.

Kamashev, D., and Rouviere-Yaniv, J. (2000) The histone-like protein HU binds specifically to DNA recombination and repair intermediates. The EMBO Journal 19: 6527–6535 10.1093/emboj/19.23.6527. Accessed June 20, 2023.

Kapanidis, A.N., Uphoff, S., and Stracy, M. (2018) Understanding Protein Mobility in Bacteria by Tracking Single Molecules. J Mol Biol 430: 4443–4455.

Kar, S., Edgar, R., and Adhya, S. (2005) Nucleoid remodeling by an altered HU protein: Reorganization of the transcription program. Proceedings of the National Academy of Sciences 102: 16397–16402 10.1073/pnas.0508032102. Accessed July 12, 2023.

Karas, V.O., Westerlaken, I., and Meyer, A.S. (2015) The DNA-Binding Protein from Starved Cells (Dps) Utilizes Dual Functions To Defend Cells against Multiple Stresses. Journal of Bacteriology 197: 3206–3215 http://www.ncbi.nlm.nih.gov/pmc/articles/PMC4560292/.

Kehrer, J.P. (2000) The Haber–Weiss reaction and mechanisms of toxicity. Toxicology 149: 43–50.

Krisko, A., and Radman, M. (2013) Biology of extreme radiation resistance: the way of Deinococcus radiodurans. Cold Spring Harbor perspectives in biology 5.

Lal, A., Dhar, A., Trostel, A., Kouzine, F., Seshasayee, A.S.N., and Adhya, S. (2016) Genome scale patterns of supercoiling in a bacterial chromosome. Nat Commun 7: 11055.

Li, S., and Waters, R. (1998) *Escherichia coli* Strains Lacking Protein HU Are UV Sensitive due to a Role for HU in Homologous Recombination. J Bacteriol 180: 3750–3756.

Liu, Y., Zhou, J., Omelchenko, M.V., Beliaev, A.S., Venkateswaran, A., Stair, J., et al. (2003) Transcriptome dynamics of *Deinococcus radiodurans* recovering from ionizing radiation. Proc Natl Acad Sci U S A 100: 4191–6. Epub 2003 Mar 21.

Macvanin, M., and Adhya, S. (2012) Architectural organization in *E. coli* nucleoid. Biochimica et Biophysica Acta (BBA) - Gene Regulatory Mechanisms 1819: 830–835.

Makarova, K.S., Aravind, L., Wolf, Y.I., Tatusov, R.L., Minton, K.W., Koonin, E.V., and Daly, M.J. (2001) Genome of the extremely radiation-resistant bacterium *Deinococcus radiodurans* viewed from the perspective of comparative genomics. Microbiol Mol Biol Rev 65: 44–79.

Mennecier, S., Coste, G., Servant, P., Bailone, A., and Sommer, S. (2004) Mismatch repair ensures fidelity of replication and recombination in the radioresistant organism *Deinococcus radiodurans*. Mol Genet Genomics 272: 460–9.

Meyer, A.S., and Grainger, D.C. (2013) The Escherichia coli Nucleoid in Stationary Phase. In Advances in Applied Microbiology. Elsevier, pp. 69–86 https://linkinghub.elsevier.com/retrieve/pii/B9780124076785000027. Accessed September 2, 2022.

Miangolarra, A.M., Li, S.H.-J., Joanny, J.-F., Wingreen, N.S., and Castellana, M. (2021) Steric interactions and out-of-equilibrium processes control the internal organization of bacteria. Proceedings of the National Academy of Sciences 118: e2106014118 10.1073/pnas.2106014118. Accessed July 13, 2023.

Mitosch, K., Rieckh, G., and Bollenbach, T. (2019) Temporal order and precision of complex stress responses in individual bacteria. Molecular Systems Biology 15: e8470 10.15252/msb.20188470. Accessed July 12, 2023.

Morikawa, K., Ushijima, Y., Ohniwa, R.L., Miyakoshi, M., and Takeyasu, K. (2019) What Happens in the Staphylococcal Nucleoid under Oxidative Stress? Microorganisms 7: 631.

Nair, S., and Finkel, S.E. (2004) Dps protects cells against multiple stresses during stationary phase. J Bacteriol 186: 4192–4198.

Nguyen, H.H., Tour, C.B. de la, Toueille, M., Vannier, F., Sommer, S., and Servant, P. (2009) The essential histone-like protein HU plays a major role in *Deinococcus radiodurans* nucleoid compaction. Molecular Microbiology 73: 240–52.

Oberto, J., Nabti, S., Jooste, V., Mignot, H., and Rouviere-Yaniv, J. (2009) The HU Regulon Is Composed of Genes Responding to Anaerobiosis, Acid Stress, High Osmolarity and SOS Induction. PLoS ONE 4: e4367.

Odsbu, I., and Skarstad, K. (2014) DNA compaction in the early part of the SOS response is dependent on RecN and RecA. Microbiology 160: 872–882.

Okanishi, H., Kim, K., Fukui, K., Yano, T., Kuramitsu, S., and Masui, R. (2017) Proteome-wide identification of lysine succinylation in thermophilic and mesophilic bacteria. Biochimica et Biophysica Acta (BBA) - Proteins and Proteomics 1865: 232–242.

Ovesný, M., Křížek, P., Borkovec, J., Švindrych, Z., and Hagen, G.M. (2014) ThunderSTORM: a comprehensive ImageJ plug-in for PALM and STORM data analysis and super-resolution imaging. Bioinformatics 30: 2389– 2390 10.1093/bioinformatics/btu202. Accessed June 30, 2023.

Passot, F.M., Nguyen, H.H., Dard-Dascot, C., Thermes, C., Servant, P., Espeli, O., and Sommer, S. (2015) Nucleoid organization in the radioresistant bacterium *Deinococcus radiodurans*. Mol Microbiol 97: 759–74.

Piggot, P.J., and Hilbert, D.W. (2004) Sporulation of *Bacillus subtilis*. Curr Opin Microbiol 7: 579–586.

Remesh, S.G., Verma, S.C., Chen, J.-H., Ekman, A.A., Larabell, C.A., Adhya, S., and Hammel, M. (2020) Nucleoid remodeling during environmental adaptation is regulated by HU-dependent DNA bundling. Nature Communications 11: 2905 10.1038/s41467-020-16724-5.

Richa, Sinha, R.P., and H Ä Der, D.P. (2015) Physiological aspects of UV-excitation of DNA. Topics in Current Chemistry 356: 203–248.

Romao, C.V., Mitchell, E.P., and McSweeney, S. (2006) The crystal structure of *Deinococcus radiodurans* Dps protein (DR2263) reveals the presence of a novel metal centre in the N terminus. J Biol Inorg Chem 11: 891– 902.

Santos, S.P., Mitchell, E.P., Franquelim, H.G., Castanho, M.A., Abreu, I.A., and Romao, C.V. (2015) Dps from *Deinococcus radiodurans*: oligomeric forms of Dps1 with distinct cellular functions and Dps2 involved in metal storage. Febs J 282: 4307–27.

Santos, S.P., Yang, Y., Rosa, M.T.G., Rodrigues, M.A.A., De La Tour, C.B., Sommer, S., et al. (2019) The interplay between Mn and Fe in *Deinococcus radiodurans* triggers cellular protection during paraquat-induced oxidative stress. Sci Rep 9: 17217.

Sato, Y.T., Watanabe, S., Kenmotsu, T., Ichikawa, M., Yoshikawa, Y., Teramoto, J., et al. (2013) Structural Change of DNA Induced by Nucleoid Proteins: Growth Phase-Specific Fis and Stationary Phase-Specific Dps. Biophysical Journal 105: 1037–1044 10.1016/j.bpj.2013.07.025. Accessed March 3, 2022.

Schneider, C.A., Rasband, W.S., and Eliceiri, K.W. (2012) NIH Image to ImageJ: 25 years of image analysis. Nature Methods 9: 671–675 10.1038/nmeth.2089.

Shechter, N., Zaltzman, L., Weiner, A., Brumfeld, V., Shimoni, E., Fridmann-Sirkis, Y., and Minsky, A. (2013) Stress-induced Condensation of Bacterial Genomes Results in Re-pairing of Sister Chromosomes. Journal of Biological Chemistry 288: 25659–25667.

Slade, D., and Radman, M. (2011) Oxidative stress resistance in *Deinococcus radiodurans*. Microbiol Mol Biol Rev 75: 133–91.

Sobetzko, P., Travers, A., and Muskhelishvili, G. (2012) Gene order and chromosome dynamics coordinate spatiotemporal gene expression during the bacterial growth cycle. Proc Natl Acad Sci USA 109 https://pnas.org/doi/full/10.1073/pnas.1108229109. Accessed September 12, 2023.

Steil, L., Serrano, M., Henriques, A.O., and Völker, U. (2005) Genome-wide analysis of temporally regulated and compartment-specific gene expression in sporulating cells of *Bacillus subtilis*. Microbiology (Reading) 151: 399–420.

Stojkova, P., Spidlova, P., and Stulik, J. (2019) Nucleoid-Associated Protein HU: A Lilliputian in Gene Regulation of Bacterial Virulence. Frontiers in Cellular and Infection Microbiology 9 https://www.frontiersin.org/articles/10.3389/fcimb.2019.00159.

Stracy, M., Schweizer, J., Sherratt, D.J., Kapanidis, A.N., Uphoff, S., and Lesterlin, C. (2021) Transient non-specific DNA binding dominates the target search of bacterial DNA-binding proteins. Molecular Cell 81: 1499–1514.e6 10.1016/j.molcel.2021.01.039. Accessed June 20, 2023.

Stringer, C., Wang, T., Michaelos, M., and Pachitariu, M. (2021) Cellpose: a generalist algorithm for cellular segmentation. Nature Methods 18: 100–106 https://app.readcube.com/library/undefined/item/5d567c6e-6e34-47aa-965f-65f37acff83e.

Szafran, M.J., Jakimowicz, D., and Elliot, M.A. (2020) Compaction and control—the role of chromosome-organizing proteins in *Streptomyces*. FEMS Microbiology Reviews 44: 725–739.

Timmins, J., and Moe, E. (2016) A decade of biochemical and structural studies of the DNA repair machinery of *Deinococcus radiodurans*. Computational and Structural Biotechnology Journa 1l 4: 168–176.

Verma, S.C., Harned, A., Narayan, K., and Adhya, S. (2023) Non-specific and specific DNA binding modes of bacterial histone, HU, separately regulate distinct physiological processes through different mechanisms. Molecular Microbiology 119: 439–455 10.1111/mmi.15033. Accessed June 20, 2023.

Verma, S.C., Qian, Z., and Adhya, S.L. (2019) Architecture of the *Escherichia coli* nucleoid. PLoS Genet 15: e1008456.

Woldringh, C.L., Jensen, P.R., and Westerhoff, H.V. (1995) Structure and partitioning of bacterial DNA: determined by a balance of compaction and expansion forces? FEMS Microbiol Lett 131: 235–242.

Zahradka, K., Slade, D., Bailone, A., Sommer, S., Averbeck, D., Petranovic, M., et al. (2006) Reassembly of shattered chromosomes in *Deinococcus radiodurans*. Nature 443: 569–73.

Zimmerman, J.M., and Battista, R.J. (2005) A ring-like nucleoid is not necessary for radioresistance in the *Deinococcaceae*. BMC Microbiology 5.

