## Supplementary information for "Stress-induced nucleoid remodeling in *Deinococcus radiodurans* is associated with major changes in HU abundance and dynamics"

**Supplementary Material**

Supplementary Discussion of the sptPALM data
Supplementary Methods
Supplementary references
Supplementary Tables S1-S3
Supplementary Figures S1-S15

------------------------------------------------------------------------------------------------------------------

**Supplementary Discussion of the sptPALM data**

sptPALM can provide detailed insight into the diffusion behavior of molecules. However, the analysis of sptPALM data is complicated by confinement (*e.g* imposed by the cell or nucleoid shape), which is particularly true in bacteria because of their small size. Confinement has been reported to affect sptPALM in two ways: (i) it reduces the apparent diffusion coefficient of molecules (Stracy *et al.*, 2021; Śmigiel *et al.*, 2022) and (ii) it creates a spatial heterogeneity in the apparent diffusion coefficient of molecules (Śmigiel *et al.*, 2022). Using the SMIS simulations software (Bourgeois, 2023), we assessed how the apparent diffusion of HU-mEos4b is affected by its confinement in *D. radiodurans* nucleoids and whether, despite the confinement, it is possible to extract the number of diffusive populations, their relative sizes and (apparent) diffusion coefficients from the diffusion coefficient histograms.

Given that confinement decreases the apparent diffusion coefficient, we first investigated whether the apparent reduction in diffusion of HU-mEos4b in stationary phase cells could be explained by an increased level of confinement due to a decreased nucleoid volume upon entry into stationary phase. To test this hypothesis, diffusion of molecules (D=0.3 $\mu m^2/s$) was simulated in 3D inside segmented nucleoids ($DR^{WT}$ syto9 staining) of exponential and stationary phase cells (Figure 1A and 1B). The diffusion coefficient used for the simulations was chosen so that the resulting apparent diffusion coefficient was in the same range as the apparent diffusion coefficient of HU-mEos4b determined experimentally in exponential phase *D. radiodurans* cells. The simulations revealed that the apparent diffusion coefficient of molecules diffusing with D = 0.3 $\mu m^2/s$ is not considerably lower in segmented stationary phase nucleoids than in exponential phase nucleoids (Figure 1C). This simulation, thus, indicates that increased confinement is insufficient to explain the decreased diffusion of HU-mEos4b upon entry into stationary phase, suggesting that the reduction in diffusion is caused by a change in the interaction between HU-mEos4b and the DNA.

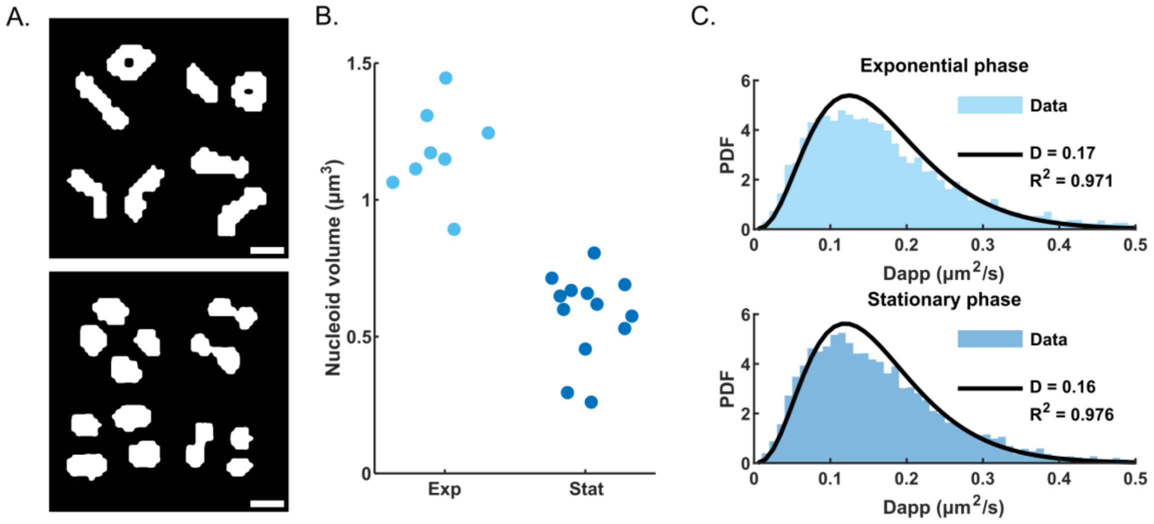


**Figure 1. Confinement alone is insufficient to explain the apparent reduction in diffusion of HU-mEos4b in stationary phase cells.** A) Z-slices of the segmented nucleoid volumes used for the simulations. Top: nucleoids of exponential phase cells; Bottom: nucleoids of stationary phase cells. Scale bar = 1 µm. B) Volume of the exponential (Exp, light blue) and stationary (Stat, dark blue) phase segmented nucleoids used for the simulations. C) Histograms of the apparent diffusion coefficient ($D_{app}$) of simulated molecules (D=0.3 µm$^2$/s) diffusing in the nucleoid volumes of exponential (top, light blue) and stationary (bottom, dark blue) phase cells (shown in A/B). The black lines show a fit of the data to a 1-population model. $R^2$ values for the fits and the derived $D_{app}$ values (in µm$^2$/s) are indicated in each case.

Although both of the simulations shown in Figure 1C contained only a single diffusive population, the $D_{app}$ distributions were not well described by a 1-population fit. This is likely explained by the heterogeneity in the $D_{app}$ created by confinement (Śmigiel *et al.*, 2022), which might be aggravated in *D. radiodurans* nucleoids by the natural variation in nucleoid shapes (and volumes) (as seen in Figure 1 A/B). Indeed, while a 1-population model cannot describe the $D_{app}$ distributions of simulated molecules confined in *D. radiodurans* nucleoid volumes, it can describe the $D_{app}$ distribution of unconfined molecules (Figure 2A). The effect of confinement can also be visualized by a spatial map of the average jump distance of the simulated data (Figure 2B), which shows that diffusion appears slower at the borders of the confining volume than in the center. We examined whether this 'border effect' is also present in the experimental data and found that this is indeed the case (Figure 2C), although the maps are noisier.

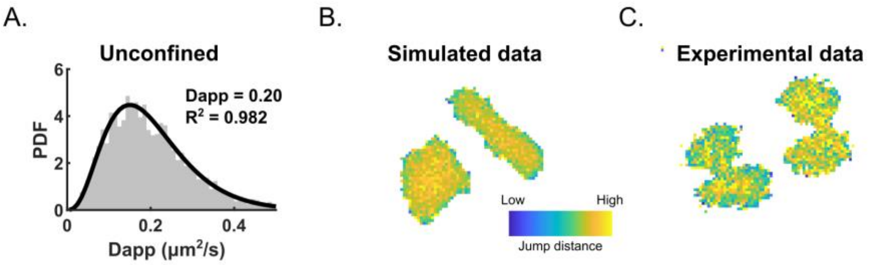


**Figure 2. Confinement creates a heterogeneity in the apparent diffusion of molecules.** A) $D_{app}$ histogram of simulated molecules (D=0.3 µm²/s) diffusing without confinement. B) Diffusion map showing the relative mean jump distance (pixel size = 50 nm) of simulated molecules confined in exponential phase nucleoid volumes. C) Diffusion map as in (B) of experimental data (exponential phase).

This distortion of the $D_{app}$ distribution due to confinement poses a problem for the fitting of the distribution in view of extracting the number of diffusive populations and their corresponding sizes and diffusion coefficients. The simulated data (see Figure 1, D = 0.3 µm²/s) cannot be described by a 1-population model, but are well fitted by a 2-population model (Figure 3A). The same is true for the experimental data (Figure 3B). This suggests that we cannot reliably determine the number of diffusive populations present in our data (1 or more). In the literature, the diffusion of DNA-binding proteins, including HU, is often described by a 2-population model (Bettridge *et al.*, 2021; Stracy *et al.*, 2021), but single diffusive species have also been reported (Floc'h *et al.*, 2019). Therefore, given the distortion due to confinement and the uncertainty in the literature, we chose not to fit the sptPALM data presented in this study.

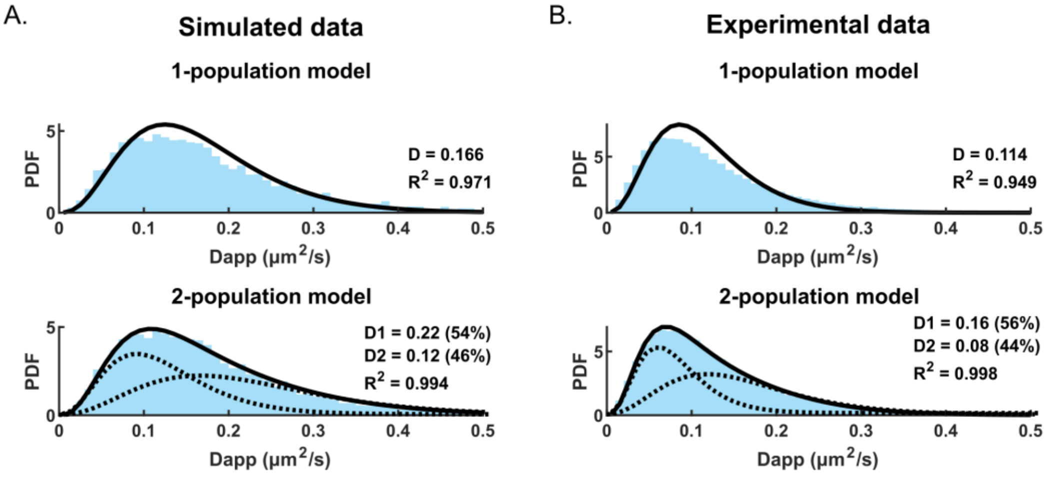


**Figure 3. Confinement hinders fitting of the $D_{app}$ distributions.** Fitting of the $D_{app}$ distributions of simulated molecules confined in exponential phase *D. radiodurans* nucleoid volumes (A, see Figure 1) and of experimental data (B, exponential phase) with a 1-population model (top row) and 2-population model (bottom row). $R^2$ values

for the fits and the derived $D_{app}$ values (in µm$^2$/s) are indicated in each case. Solid lines depict the fit; dashed lines depict the individual populations of the 2-population fit.

**Supplementary Methods**

sptPALM simulations were performed using the recently developed software SMIS (Bourgeois, 2023). Experimentally obtained masks of *D. radiodurans* nucleoids in exponential and stationary phase ($DR^{WT}$ syto9 staining) were used as confining volumes. Simulations were performed in 3D using a frame time of 10 ms. The simulated data were processed similarly as experimental data using Thunderstorm (Ovesný *et al.*, 2014), SWIFT (Endesfelder *et al.*, in prep) and Matlab.

The following models were used for the fitting of the $D_{app}$ ($D_i$*) distributions. The probability of measuring $D_i$* for a molecule that diffuses with an apparent diffusion coefficient D* is given by (Stracy *et al.*, 2021):

$$p(D_i^*) = \frac{1}{(n-1)!} * \left(\frac{n}{D^*}\right)^n * (D_i^*)^{n-1} * exp\left(\frac{-nD_i^*}{D^*}\right)$$

This model can be extended to include two populations with apparent diffusion coefficients $D_1^*$ and $D_2^*$ and relative populations of $A_1$ and $(1 - A_1)$:

$$p(D_i^*) = \left[\frac{A_1}{(n-1)!} * \left(\frac{n}{D_1^*}\right)^n * (D_i^*)^{n-1} * exp\left(\frac{-nD_i^*}{D_1^*}\right)\right] + \left[\frac{(1-A_1)}{(n-1)!} * \left(\frac{n}{D_2^*}\right)^n * (D_i^*)^{n-1} * exp\left(\frac{-nD_i^*}{D_2^*}\right)\right]$$

Distributions of the $D_{app}$ were fitted in MATLAB using the 1- or 2-population model using nonlinear least squares curve fitting.

**Supplementary references**

Bettridge, K., Verma, S., Weng, X., Adhya, S., and Xiao, J. (2021) Single-molecule tracking reveals that the nucleoid-associated protein HU plays a dual role in maintaining proper nucleoid volume through differential interactions with chromosomal DNA. *Molecular Microbiology* **115**: 12–27 https://doi.org/10.1111/mmi.14572. Accessed June 20, 2023.

Bourgeois, D. (2023) Single molecule imaging simulations with advanced fluorophore photophysics. *Communications Biology* **6**: 53 https://doi.org/10.1038/s42003-023-04432-x.

Endesfelder, M., Schließl, C., Turkowyd, B., Lechner, T., and Endesfelder, U. swift – fast, probabilistic tracking for dense, highly dynamic single-molecule data. **Manuscript in prep.**

Floc'h, K., Lacroix, F., Servant, P., Wong, Y.-S., Kleman, J.-P., Bourgeois, D., and Timmins, J. (2019) Cell morphology and nucleoid dynamics in dividing *Deinococcus radiodurans*. *Nature Communications* **10**: 3815 https://doi.org/10.1038/s41467-019-11725-5.

Ovesný, M., Křížek, P., Borkovec, J., Švindrych, Z., and Hagen, G.M. (2014) ThunderSTORM: a comprehensive ImageJ plug-in for PALM and STORM data analysis and super-resolution imaging. *Bioinformatics* **30**: 2389–2390 https://doi.org/10.1093/bioinformatics/btu202. Accessed June 30, 2023.

Passot, F.M., Nguyen, H.H., Dard-Dascot, C., Thermes, C., Servant, P., Espeli, O., and Sommer, S. (2015) Nucleoid organization in the radioresistant bacterium *Deinococcus radiodurans*. *Mol Microbiol* **97**: 759–74.

Śmigiel, W.M., Mantovanelli, L., Linnik, D.S., Punter, M., Silberberg, J., Xiang, L., *et al.* (2022) Protein diffusion in Escherichia coli cytoplasm scales with the mass of the complexes and is location dependent. *Science Advances* **8**: eabo5387 https://doi.org/10.1126/sciadv.abo5387. Accessed July 12, 2023.

Stracy, M., Schweizer, J., Sherratt, D.J., Kapanidis, A.N., Uphoff, S., and Lesterlin, C. (2021) Transient non-specific DNA binding dominates the target search of bacterial DNA-binding proteins. *Molecular Cell* **81**: 1499-1514.e6 https://doi.org/10.1016/j.molcel.2021.01.039. Accessed June 20, 2023.

**Supplementary Tables S1-S3**

***Table S1: Effects of sublethal and lethal UV-C on* D. radiodurans *cell and nucleoid morphology***

| STAGES | PROCESS | SUBLETHAL UV-C DOSE | LETHAL UV-C DOSE |
|---|---|---|---|
| **Stage 1** | **Cell growth & morphology** | - Septation is arrested leading to the accumulation of P3 cells.<br>- Splitting of tetrads into diads (P6->P1) is unaffected.<br>- Cell size remains constant.<br>- No visible defects in the membrane. | - Cell cycle is fully arrested.<br>- Cell size remains constant.<br>- No visible defects in the membrane. |
| | **Nucleoid size & morphology** | - Minimal nucleoid volume observed at $t_0$, which remains stable during stage 1.<br>- 80% of cells exhibit rounded nucleoids at $t_{1h}$. | - Reduced nucleoid volume observed at $t_0$, which remains stable during stage 1.<br>- 95% of cells exhibit rounded nucleoids at $t_{2h}$ with a very high sphericity value. |
| | **HU level & dynamics** | - HU abundance and diffusion rapidly increase after irradiation ($t_0$). | - HU abundance and diffusion rapidly increase after irradiation ($t_0$). |
| **Stage 2** | **Cell growth & morphology** | - Cell growth & division are partially restored.<br>- Mean cell size increases.<br>- Very large cells start to appear with visible membrane defects (up to 10-15% cells). | - Cell cycle remains largely arrested.<br>- A few surviving cells start dividing again at the end of stage 2. |
| | **Nucleoid size & morphology** | - Nucleoid volume rapidly increases again.<br>- A fraction of cells exhibits large expanded nucleoids (volume > $3\mu m^3$).<br>- A majority of cells recover a 'normal' structured nucleoid morphology. | - Nucleoids remain rounded, but their volume progressively reduces to micro-nucleoids (<$0.3\mu m^3$).<br>- At $t_{7h}$, 60% of cells have micro-nucleoids. |
| | **HU level & dynamics** | - HU abundance decreases between $t_{1h}$ and $t_{3h}$ and then remains constant.<br>- HU diffusion progressively decreases. | - HU abundance remains constant.<br>- HU diffusion continues to increase with an enhanced cell-to-cell variation. |
| **Stage 3** | **Cell growth & morphology** | - Cell population progresses towards stationary phase leading to the accumulation of P6 and P7 cells.<br>- Mean cell size decreases.<br>- 10-15% cells still exhibit abnormal cell morphology with visible membrane defects. | - Many anucleate cells exhibit distorted cell morphologies.<br>- A small fraction of surviving cells grows and divides with no obvious defects. |
| | **Nucleoid size & morphology** | - Nucleoid volume decreases – a few cells still harbor expanded nucleoids.<br>- 90% nucleoids are well structured. | - At $t_{24h}$, 95% of cells show no or diffuse Syto9 staining.<br>- Surviving cells exhibit either normal or expanded nucleoid morphologies. |
| | **HU level & dynamics** | - HU abundance increases.<br>- 'Normal' HU dynamics are largely restored. | - At $t_{24h}$, many cells exhibit very high or very low HU diffusion.<br>- HU abundance is slightly increased at $t_{24h}$. |

***Table S2: Description of strains used in this study***

| ***Strains*** | ***Description*** | ***Source or reference*** |
|---|---|---|
| ***E. coli DH5a*** | *fhuA2Δ(argF-lacZ)U169 phoA glnV44 Φ80Δ(lacZ)M15 gyrA96 recA1 relA1 endA1 thi-1 hsdR17* | *New England Biolabs* |
| ***D. radiodurans*** | | |
| ***GY9613*** | *ATCC 13939, R1* | *Laboratory stock* |
| ***GY15743*** | *hbs::cherryΩcat* | *Laboratory stock* |
| ***GY15787*** | *hbs::cherryΩtet, locus ori::kan-parSc2$_{Bc}$/pFAP246 ($P_{spac}$-parBc2Bc::drGFP, cat)* | (Passot *et al.*, 2015) |
| ***GY15800*** | *hbs::cherryΩtet, locus ter::kan-parSc2$_{Bc}$/pFAP246 ($P_{spac}$-parBc2Bc::drGFP, cat)* | (Passot *et al.*, 2015) |
| ***GY16993*** | *hbs::mEos4b-NCherryΩhph* | *This work* |

***Table S3: Oligonucleotides used to prepare the DR$^{HUmEos4b}$ strain***

| **Name** | **Sequence** |
|---|---|
| HU_1 | GTCATGCTGCTCACCATGAC |
| HU_2_BamHI | GATTGGATCCCAGGTTGCCCTTGAGGGTGC |
| HU_3_XbaI | GGAATCTAGAGCGGGTTCGTCCCTGACCGC |
| HU_4 | TCGCCGAGATGATCCTCGAC |
| mEos4b_BamHI | GCATGGATCCATGGTCTCGAAAGGGGAGGAA |
| Hph_XbaI | GGAATCTAGATTATTCCTTTGCCCTCGGACG |

### Figure S1

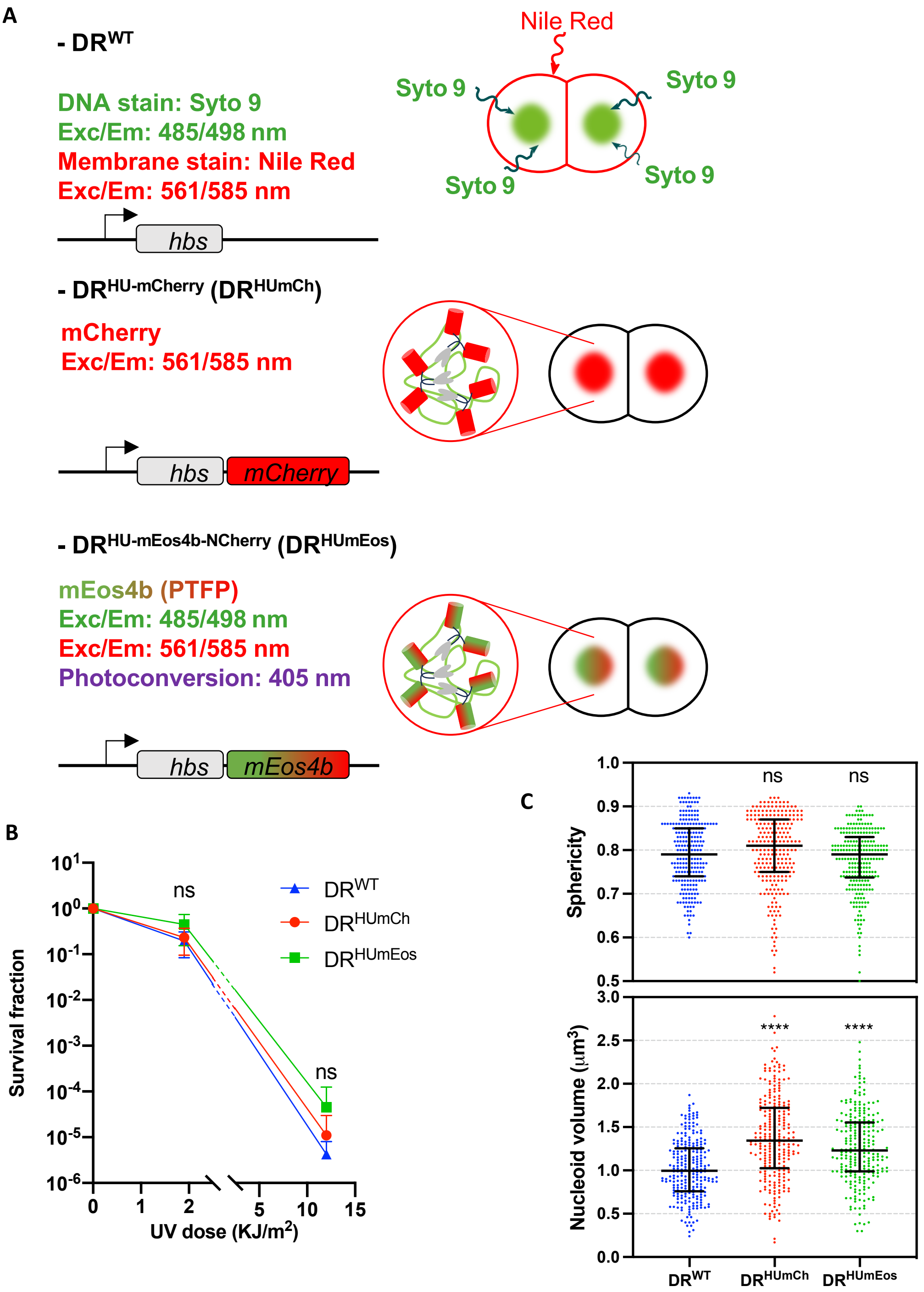


***Figure S1: Characteristics of the three strains of*** **Deinococcus radiodurans** ***used in this study.*** *(A) Nile Red and Syto9 dyes were used with WT* D. radiodurans *($DR^{WT}$) to label the plasma membrane and DNA respectively in red and green. Two genetically engineered strains expressing HU (encoded by the hbs gene) from its endogenous promoter fused to either mCherry (middle; $DR^{HUmCh}$) or mEos4b-NCherry (lower panel; $DR^{HUmEos}$; see details in Fig. S15) were also used. The excitation and emission wavelengths used for imaging these strains are indicated. (B) Survival curves of exponentially growing $DR^{WT}$ (blue), $DR^{HUmCh}$ (red) and $DR^{HUmEos}$ (green) after exposure to 1.9 (sublethal) and 12.0 (lethal) kJ/m² UV-C light. Data represent the mean and standard deviation of 3 independent experiments. (C) Nucleoid volume and sphericity of non-irradiated exponentially growing $DR^{WT}$ (blue), $DR^{HUmCh}$ (red) and $DR^{HUmEos}$ (green) cells stained with Syto9 (n=250). Error bars represent the median and interquartile range. ns: non-significant, **** $p<0.0001$, Kruskal-Wallis statistical test performed in GraphPad Prism 8.*

### Figure S2

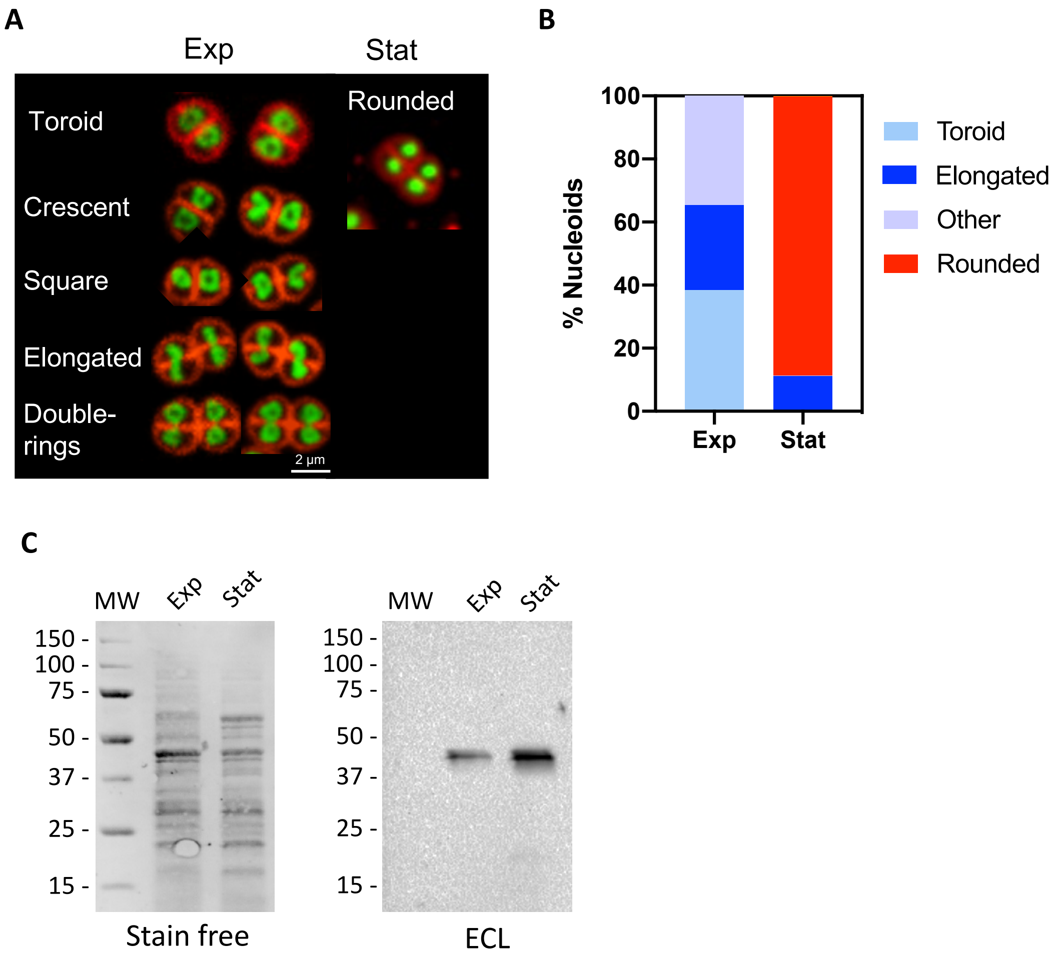


***Figure S2: Nucleoid morphology and HU expression in exponential and stationary phase*** **D. radiodurans.** *(A) Illustration of the most common 'normal' nucleoid morphologies observed in exponential and stationary phase* D. radiodurans. *(B) Distribution of the major nucleoid morphologies in exponential and stationary phase DR$^{WT}$(n>450). (C) Western blot analysis of HU-mCherry expression in exponential and stationary phase DR$^{HUmCh}$. Left: Stain free (BioRad) staining of the nitrocellulose membrane prior to incubation with the anti-mCherry antibody. This image was used to normalize the amounts of cell extract loaded in each well. Molecular weights are indicated in kDa. Right: chemiluminescent signal revealing HU-mCherry bands.*

### Figure S3

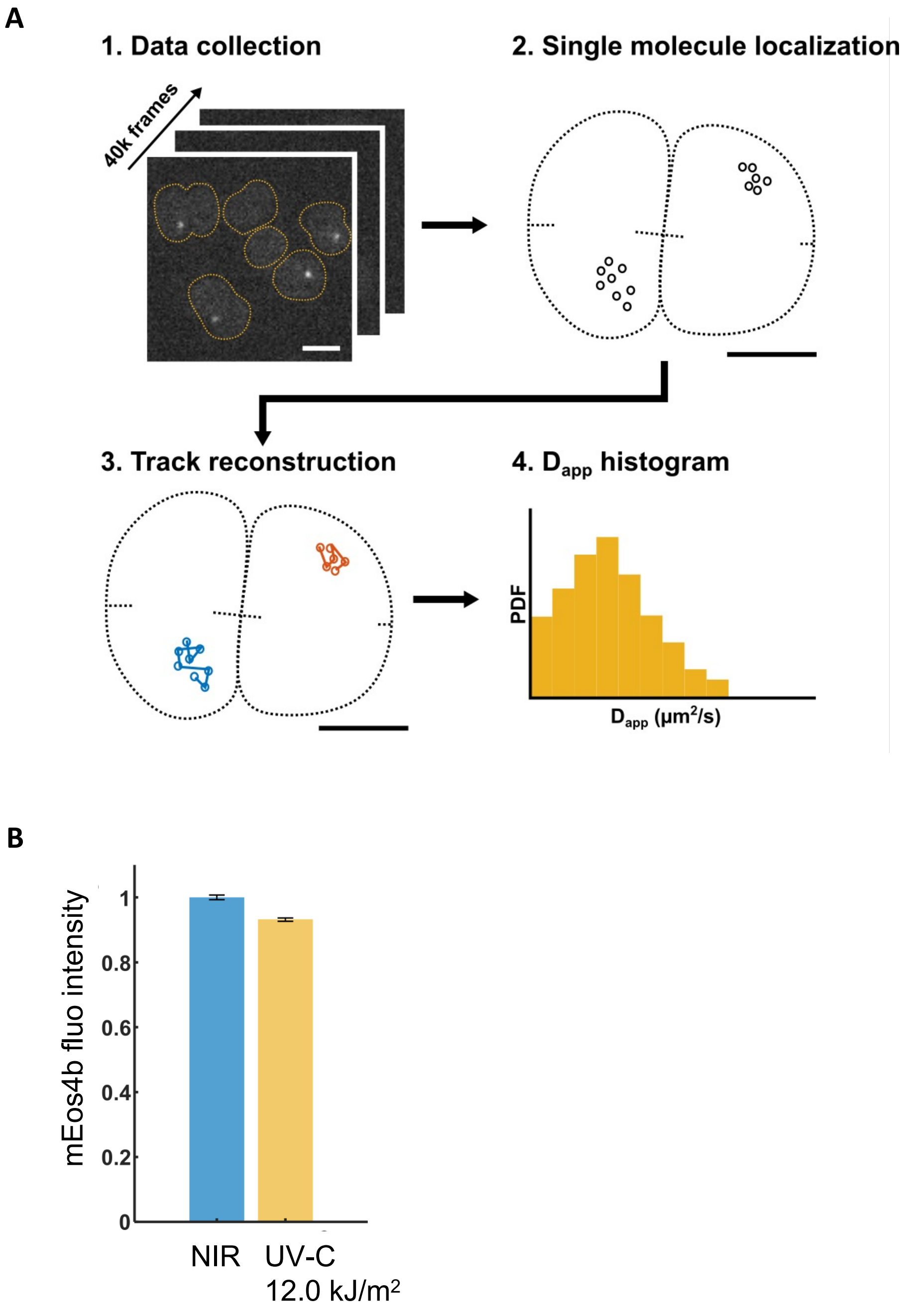


***Figure S3: Experimental set-up and data analysis pipeline used for the single-particle tracking experiments on DR$^{HUmEos}$.*** *(A) First, single-molecule data (typically 30,000-40,000 frames) were acquired on live DR$^{HUmEos}$ cells by PALM imaging (1). Next, single molecules were localized using Thunderstorm (2) and linked in time to reconstruct tracks belonging to unique molecules in SWIFT (3). Finally, the tracks were analyzed to extract the apparent diffusion coefficients ($D_{app}$) of HU-mEos4b molecules (4). PDF: Probability Density Function. (B) Comparison of the mean normalized intensity of HU-mEos4b before (blue) and after exposure to 12.0 kJ/m$^2$ (yellow).*

### Figure S4

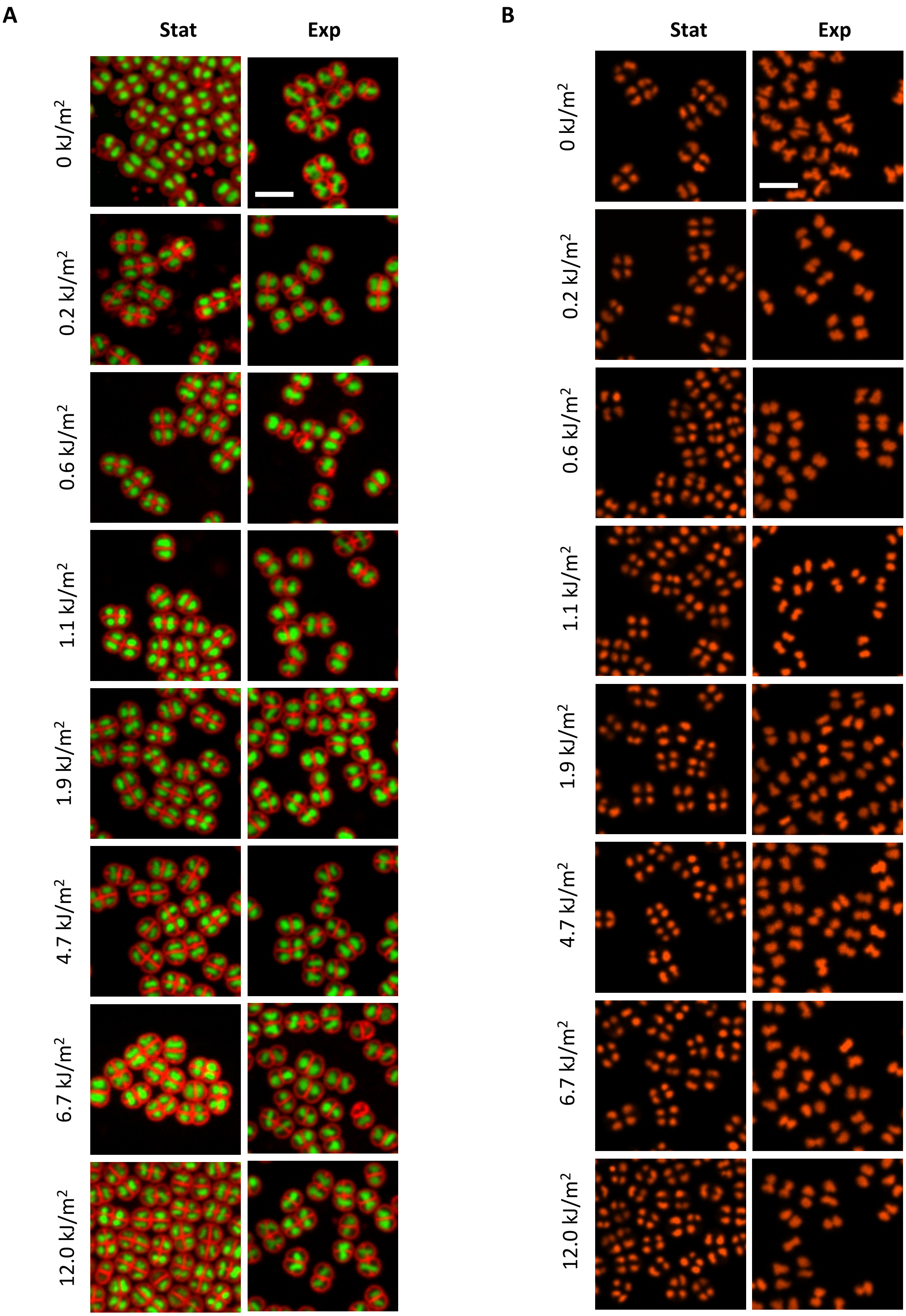


***Figure S4: Representative images of DR$^{WT}$ and DR$^{HUmCh}$ after exposure to increasing doses of UV-C light.*** *(A) Representative images of stationary (left) and exponential (right) phase DR$^{WT}$ cells stained with Nile Red (membrane) and Syto9 (nucleoids) immediately after being exposed to 0-12.0 kJ/m$^2$ UV-C light. (B) Representative images of stationary (left) and exponential (right) phase DR$^{HUmCh}$ cells immediately after being exposed to 0-12.0 kJ/m$^2$ UV-C light. Scale bar = 5 µm.*

### Figure S5

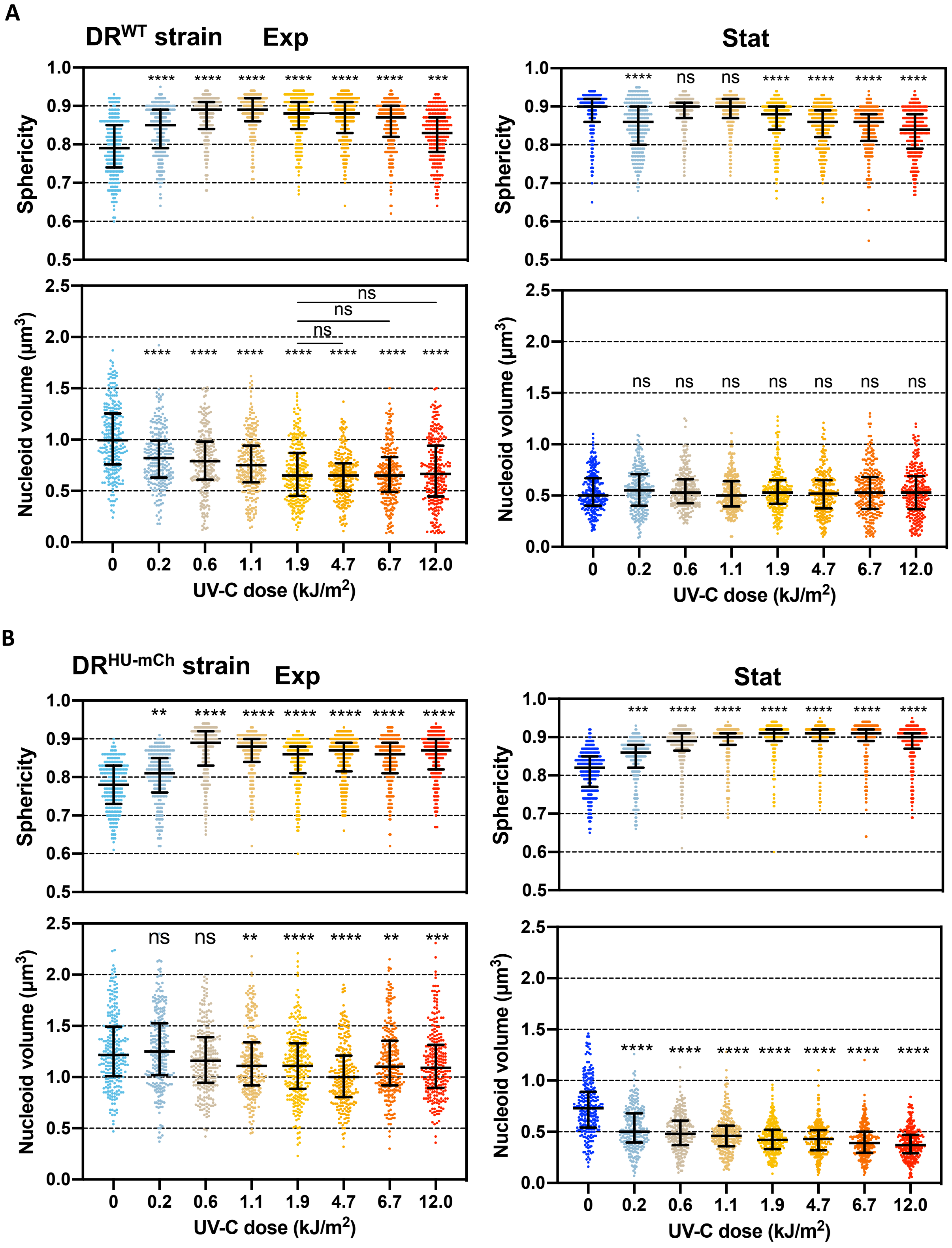


***Figure S5: Changes in nucleoid volume and sphericity as a function of UV-C dose.*** *(A)-(B) Nucleoid volume and sphericity of exponential (left) and stationary (right) phase $DR^{WT}$ (A) and $DR^{HUmCh}$ (B) after exposure to 0, 0.2, 0.6, 1.1, 1.9, 4.7, 6.7 and 12.0 $kJ/m^2$ UV-C light (n=250). Error bars represent the median and interquartile range. ns: non-significant, ** $p<0.01$, *** $p<0.001$, **** $p<0.0001$, Kruskal-Wallis statistical test performed in GraphPad Prism 8.*

### Figure S6

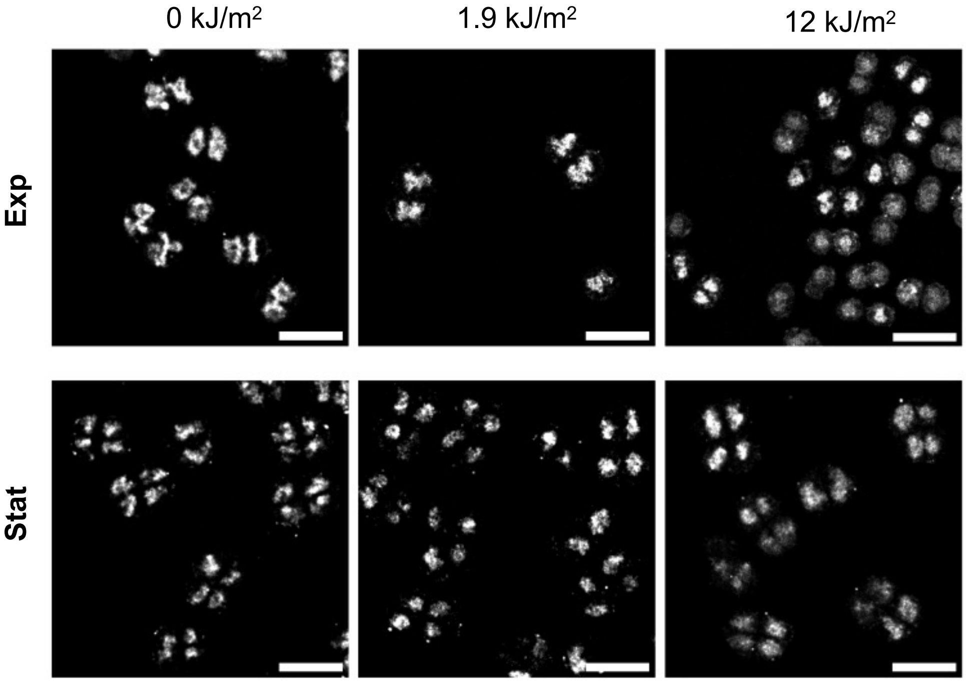


***Figure S6: Representative images of DR$^{HU\text{-}mEos}$ nucleoids before (left) and after exposure to 1.9 (middle) and 12.0 kJ/m$^2$ UV-C light (right).*** *Scale bar = 5 µm.*

### Figure S7

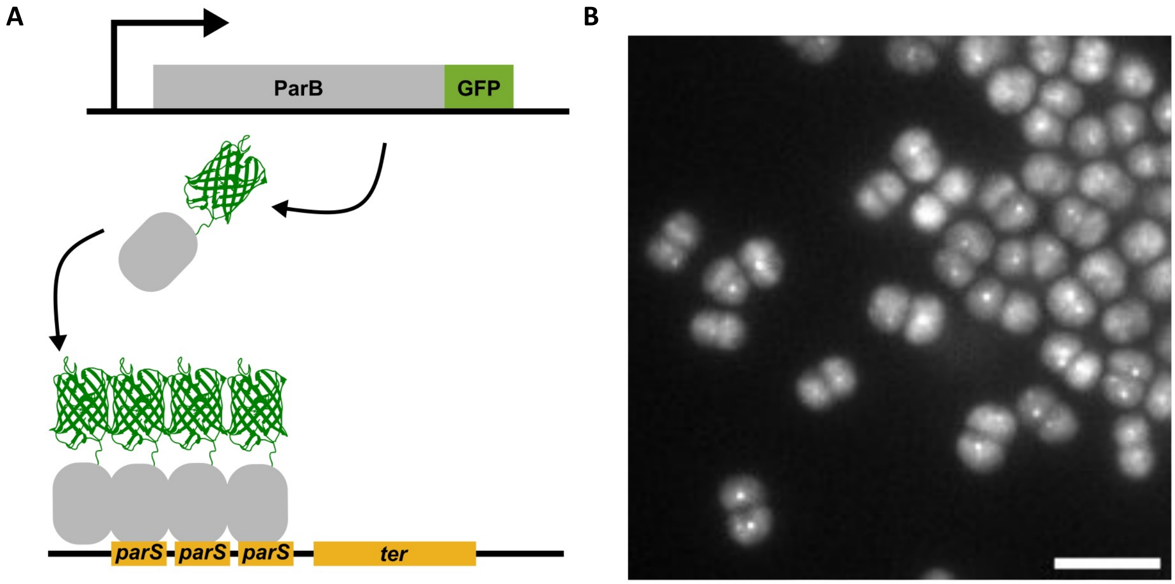


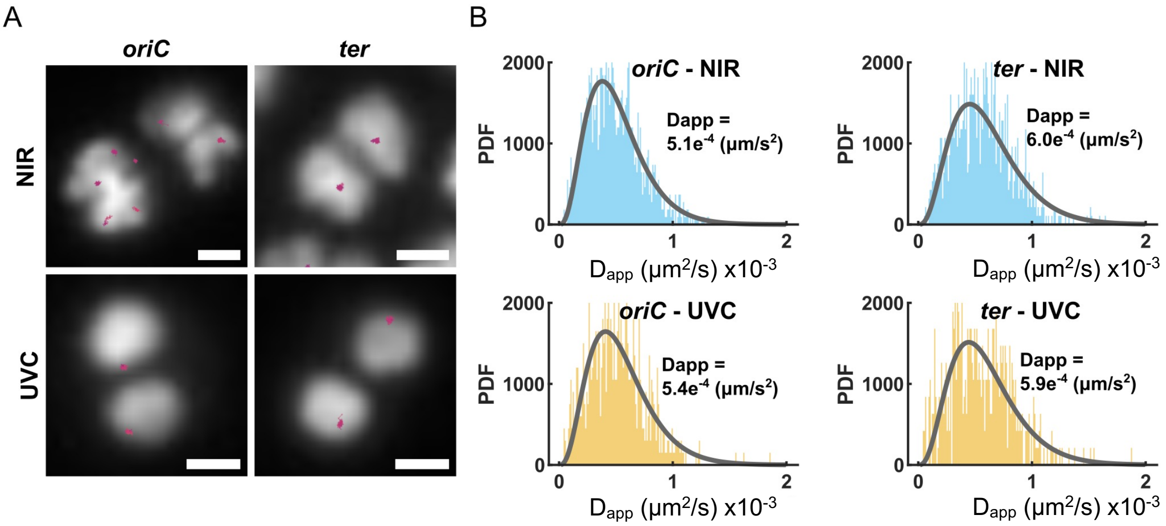


***Figure S7: Tracking of chromosome I oriC and ter foci after exposure of exponential phase* D. radiodurans *to 1.9kJ/m² UV-C light*** *(A) Schematic diagram illustrating the labeling of the ter (or oriC) locus of* D. radiodurans *chromosome I through a heterologous parS/ParB system* (Passot *et al.*, 2015). *parS sites were inserted nearby the targeted loci and the cognate ParB was expressed as a GFP fusion protein from a plasmid. These genetic modifications were inserted into a HU-mCherry expressing strain of* D. radiodurans *to allow the visualization of the nucleoid as well as the ter/oriC loci. (B) Representative image of ParB-GFP labeled ter foci in* D. radiodurans. *The high fluorescence background in the cells arises from freely diffusing ParB-GFP. Scale bar = 5 µm. (C) Representative tracks of the oriC and ter foci (purple) in non-irradiated (NIR) and UV-C irradiated (1.9 kJ/m²) cells after 1 hour of recovery overlayed on HU-mCherry labeled nucleoids (white). Scale bar = 500 nm. (D) Histogram distributions of the apparent diffusion coefficients ($D_{app}$) of the oriC (left) and ter (right) loci in non-irradiated (NIR; top plots) and irradiated cells (UVC; lower plots) after1 hour of recovery. The data were fitted with a 1-population fit and the derived $D_{app}$ values are indicated. n=475-1000 tracks from 2 experiments.*

### Figure S8

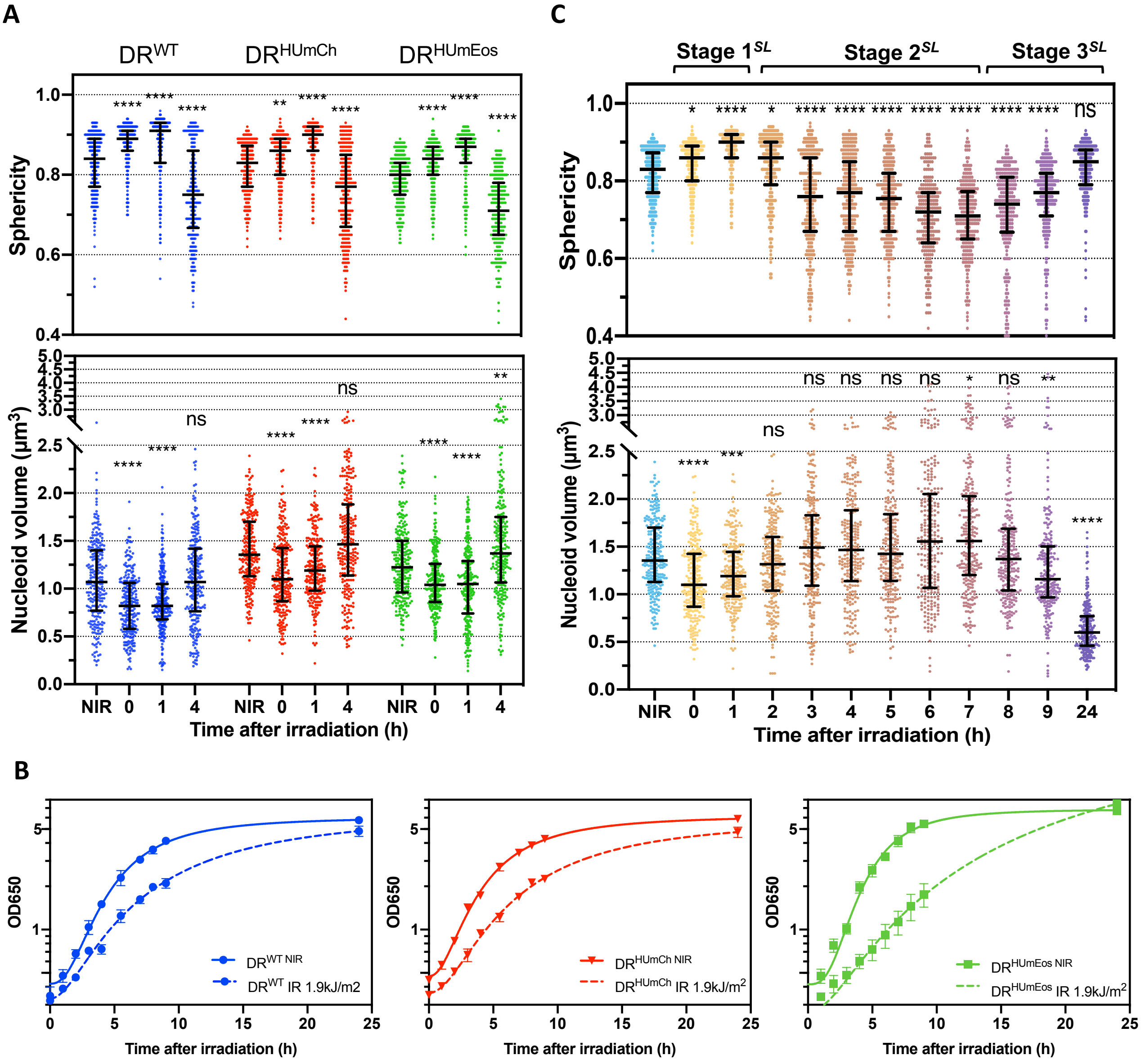


***Figure S8: Effects of sublethal UV-C light on cell growth and on nucleoid shape and size.*** *(A) Changes in nucleoid volume and sphericity in exponentially growing $DR^{WT}$ (blue), $DR^{HUmCh}$ (red) and $DR^{HUmEos}$ (green) cells following exposure to 1.9 kJ/m2 UV-C light (n=250). NIR: non-irradiated. Nucleoid volume and sphericity were measured immediately after irradiation (0), 1h and 4h post-irradiation. (B) Growth curves of non-irradiated (NIR; full line) and irradiated (1.9 $kJ/m^2$; dashed line) $DR^{WT}$ (blue), $DR^{HUmCh}$ (red) and $DR^{HUmEos}$ (green). Data represent the mean and standard deviation of 3 independent experiments. (C) Detailed analysis of the changes in nucleoid volume and sphericity in exponentially growing $DR^{HUmCh}$. Data points of non-irradiated samples are illustrated in light blue, while those corresponding to the different timepoints during the recovery phase are colored from yellow to purple. (A)-(C) Error bars represent the median and interquartile range. ns: non-significant, * p<0.05, ** p<0.01, *** p<0.001, **** p<0.0001, Kruskal-Wallis statistical test performed in GraphPad Prism 8.*

### Figure S9

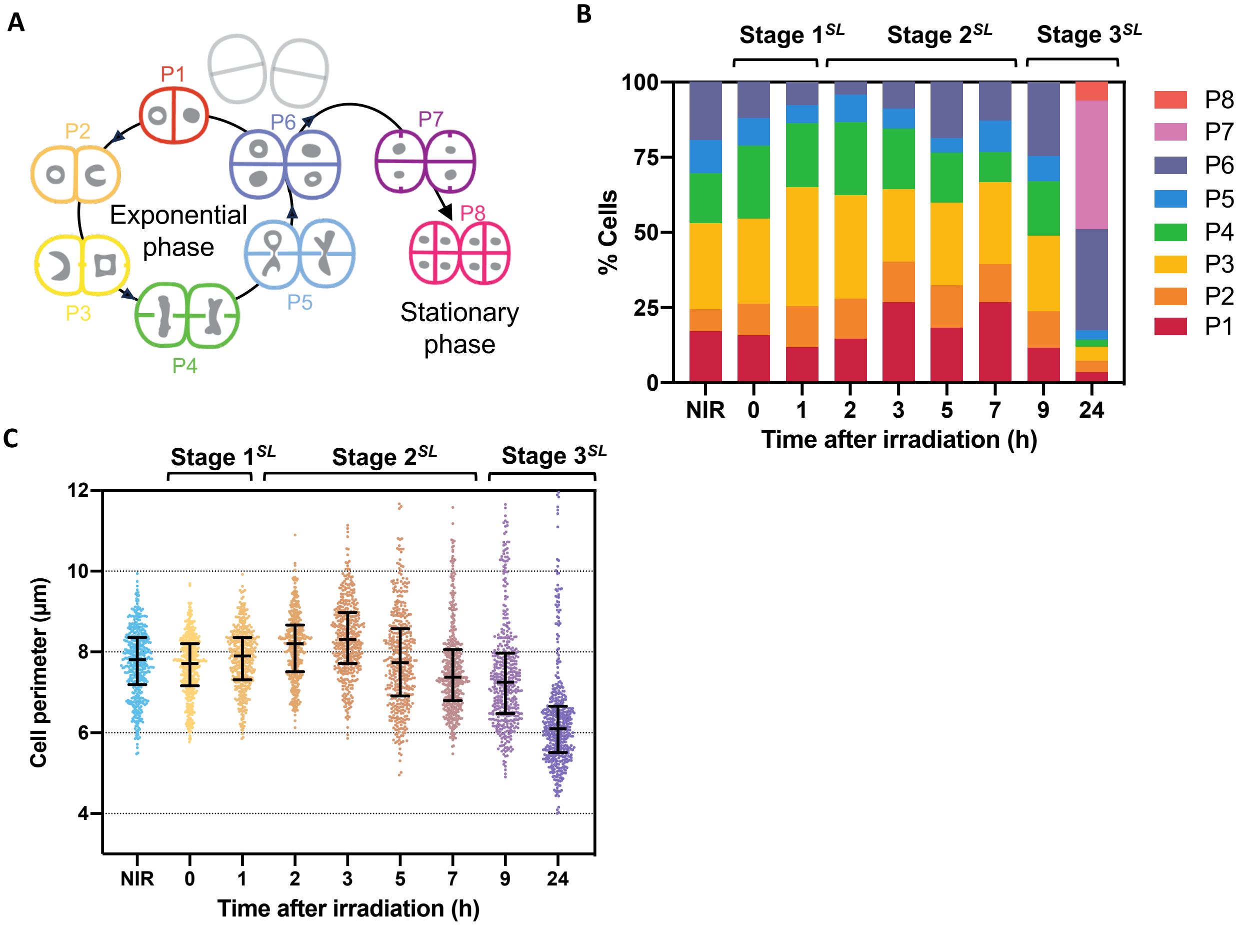


***Figure S9: Effects of sublethal UV-C light on cell growth and division in DR$^{WT}$.*** *(A) Schematic diagram of the different phases of* D. radiodurans*' cell cycle. In exponential phase, a diad in phase 1 (P1) progressively grows and divides to form tetrads in phase 6 (P6). In stationary phase, P6 cells accumulate and no longer separate into diads, but instead initiate a new cycle of division to produce phase 7 and 8 cells. (B) Distribution of cells in the various phases of the cell cycle before (NIR; non-irradiated) and after exposure to sublethal (1.9 kJ/m$^2$) UV-C light. The distribution of cells (n>450) was determined at 0, 1, 2, 3, 5, 7, 9 and 24h post-irradiation. At the 24h timepoint, a majority of cells were entering stationary phase as evidenced by the appearance of phase 7 and 8 and the large increase in the number of phase 6 cells. (C) Evolution of the size of individual cells at different timepoints after exposure to sublethal (1.9 kJ/m$^2$) UV-C light (n>450). Error bars represent the median and interquartile range.*

### Figure S10

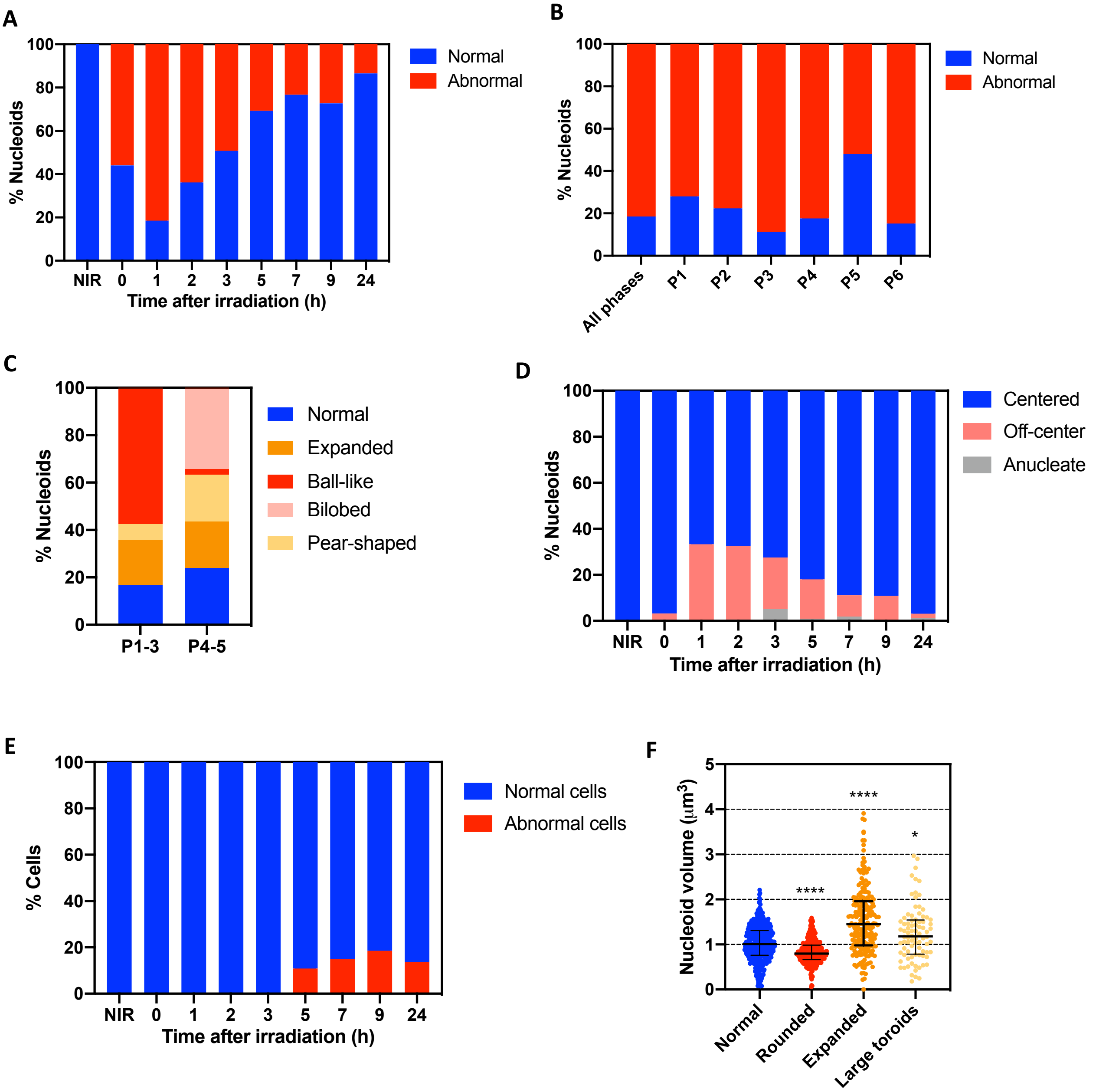


***Figure S10: Effects of sublethal UV-C light on nucleoid morphology in DR$^{WT}$.*** *(A) Percentage of normal (blue) versus abnormal (red) nucleoids at different timepoints after exposure to sublethal (1.9 kJ/m$^2$) UV-C light (n>450). (B) Percentage of normal (blue) versus abnormal (red) nucleoids at t1h after exposure to sublethal (1.9 kJ/m$^2$) UV-C light (n>450) as a function of the phase of the cell cycle. (C) Distribution of nucleoid morphologies observed for Phase 1-3 (P1-3) cells and Phase 4-5 (P4-5) cells at t1h after exposure to sublethal (1.9 kJ/m$^2$) UV-C light (n>450). (D) Percentage of centered (blue) versus off-centered (pink) nucleoids and anucleate cells (grey) at different timepoints after exposure to sublethal (1.9 kJ/m$^2$) UV-C light (n>450). (E) Percentage of normal (blue) versus abnormal (red) cells at different timepoints after exposure to sublethal (1.9 kJ/m$^2$) UV-C light (n>450). (F) Comparison of the nucleoid volumes of normal (blue), rounded (red), expanded (orange) and large toroids (yellow) (n>80). Error bars represent the median and interquartile range.*

### Figure S11

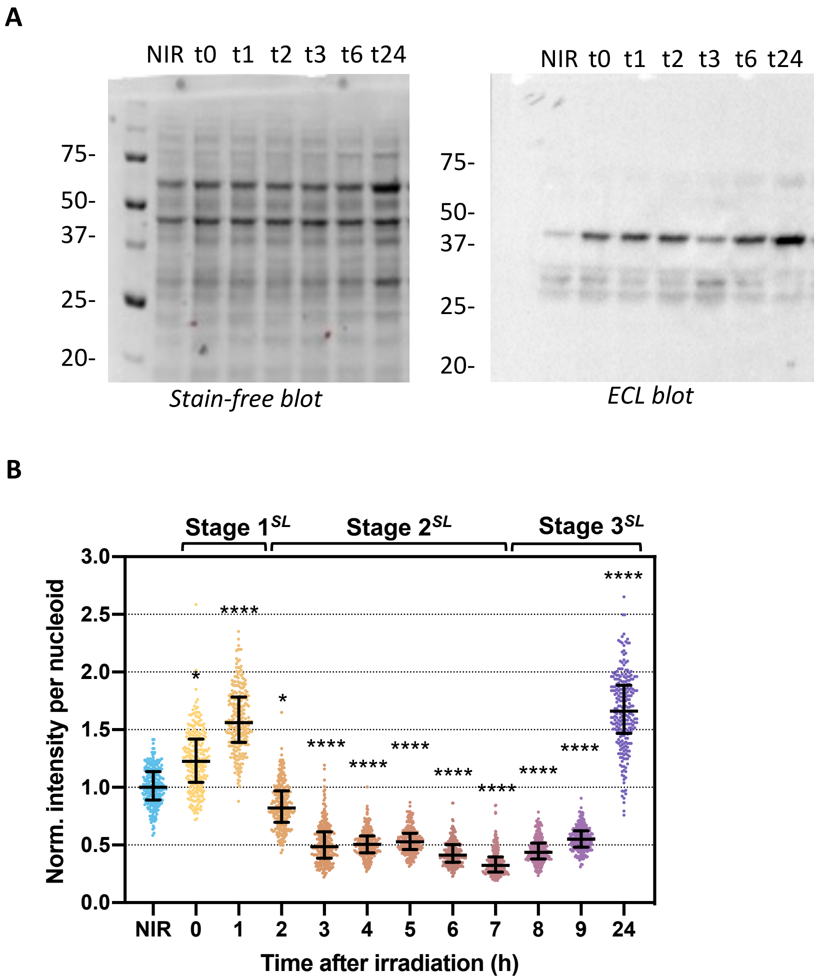


***Figure S11: HU expression after exposure to sublethal UV-C light.*** *(A) Western blot analysis of HU-mCherry expression in $DR^{HUmCh}$ at different timepoints (0, 1, 2, 3, 6 and 24h) after exposure to sublethal UV-C light. Left: Stain free (BioRad) staining of the nitrocellulose membrane prior to incubation with the anti-mCherry antibody. This image was used to normalize the amounts of cell extract loaded in each well. Molecular weights are indicated in kDa. Right: chemiluminescent signal revealing HU-mCherry bands. (B) Normalized HU-mCherry intensity in segmented nucleoids of $DR^{HUmCh}$ at different timepoints (0, 1, 2, 3, 4, 5, 6, 7, 8, 9 and 24h) after exposure to sublethal UV-C light (n>450). Error bars represent the median and interquartile range. * p<0.05, **** p<0.0001, Kruskal-Wallis statistical test performed in GraphPad Prism 8.*

### Figure S12

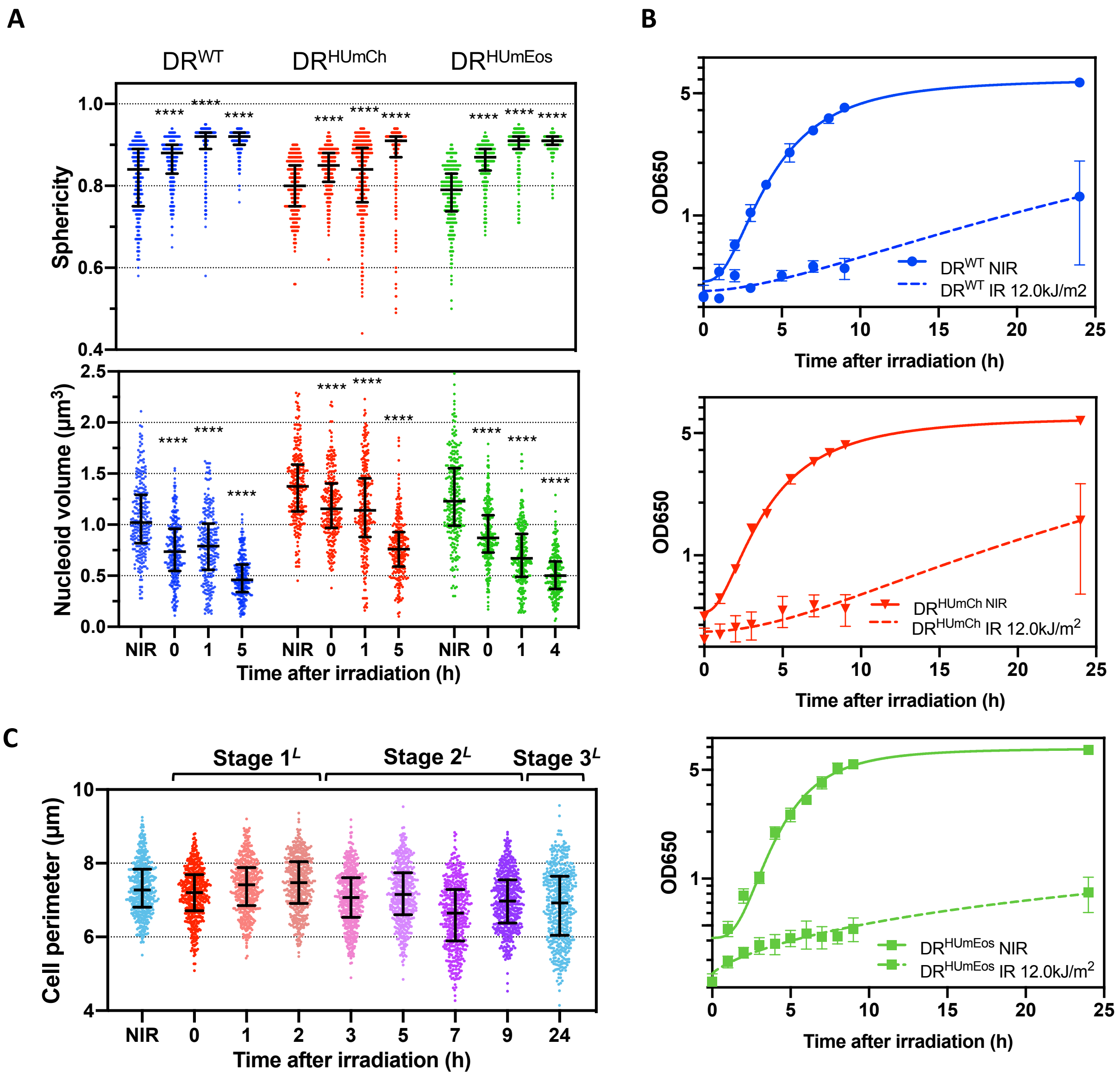


***Figure S12: Effects of lethal UV-C light on cell growth and on nucleoid shape and size.*** *(A) Changes in nucleoid volume and sphericity in exponentially growing DR$^{WT}$ (blue), DR$^{HUmCh}$ (red) and DR$^{HUmEos}$ (green) cells following exposure to 12.0 kJ/m2 UV-C light (n=250). NIR: non-irradiated. Nucleoid volume and sphericity were measured immediately after irradiation (0), 1h and 4h post-irradiation. Error bars represent the median and interquartile range. **** p<0.0001, Kruskal-Wallis statistical test performed in GraphPad Prism 8. (B) Growth curves of non-irradiated (NIR; full line) and irradiated (12.0 kJ/m$^2$; dashed line) DR$^{WT}$ (blue), DR$^{HUmCh}$ (red) and DR$^{HUmEos}$ (green). Data represent the mean and standard deviation of 3 independent experiments. (C) Evolution of the size of individual cells at different timepoints after exposure to lethal (12.0 kJ/m$^2$) UV-C light (n>450). Error bars represent the median and interquartile range. (A)-(C)*

### Figure S13

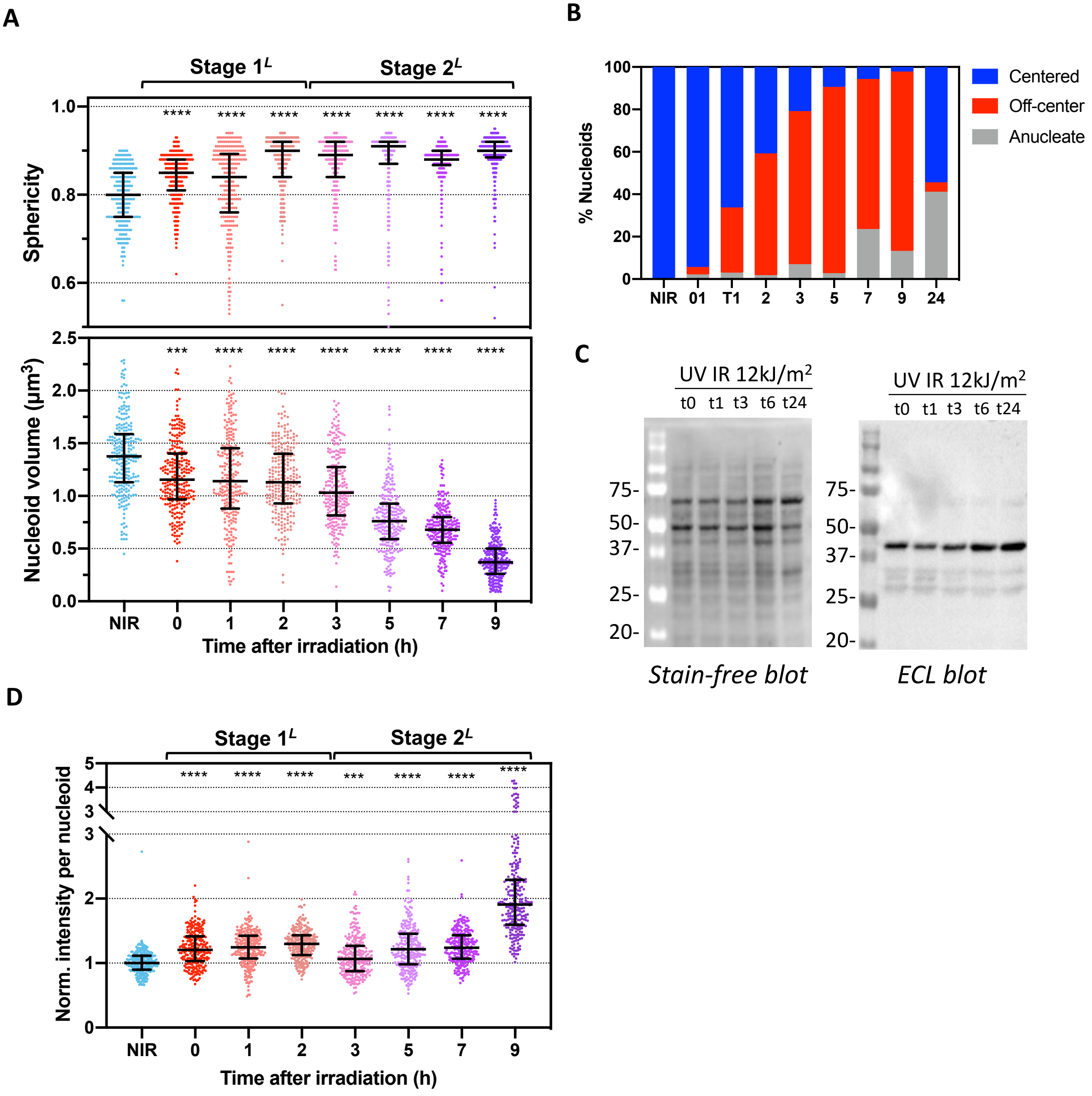


***Figure S13: Effects of lethal UV-C light on nucleoid shape and size.*** *(A) Detailed analysis of the changes in nucleoid volume and sphericity in exponentially growing DR$^{HUmCh}$. Data points for non-irradiated samples are illustrated in blue, while those corresponding to the different timepoints after exposure to lethal UV-C light are colored from red to purple. Error bars represent the median and interquartile range. *** p<0.001, **** p<0.0001, Kruskal-Wallis statistical test performed in GraphPad Prism 8. (B) Percentage of centered (blue) versus off-centered (red) nucleoids and anucleate cells (grey) at different timepoints after exposure to lethal (12.0 kJ/m²) UV-C light (n>450). (C) Western blot analysis of HU-mCherry expression in DR$^{HUmCh}$ at different timepoints (0, 1, 3, 6 and 24h) after exposure to lethal UV-C light. Left: Stain free (BioRad) staining of the nitrocellulose membrane prior to incubation with the anti-mCherry antibody. This image was used to normalize the amounts of cell extract loaded in each well. Molecular weights are indicated in kDa. Right: chemiluminescent signal revealing HU-mCherry bands. (D) Normalized HU-mCherry intensity in segmented nucleoids of DR$^{HUmCh}$ at different timepoints (0, 1, 2, 3, 5, 7, and 9h) after exposure to lethal UV-C light (n>450). Error bars represent the median and interquartile range. **** p<0.0001, Kruskal-Wallis statistical test performed in GraphPad Prism 8.*

### Figure S14

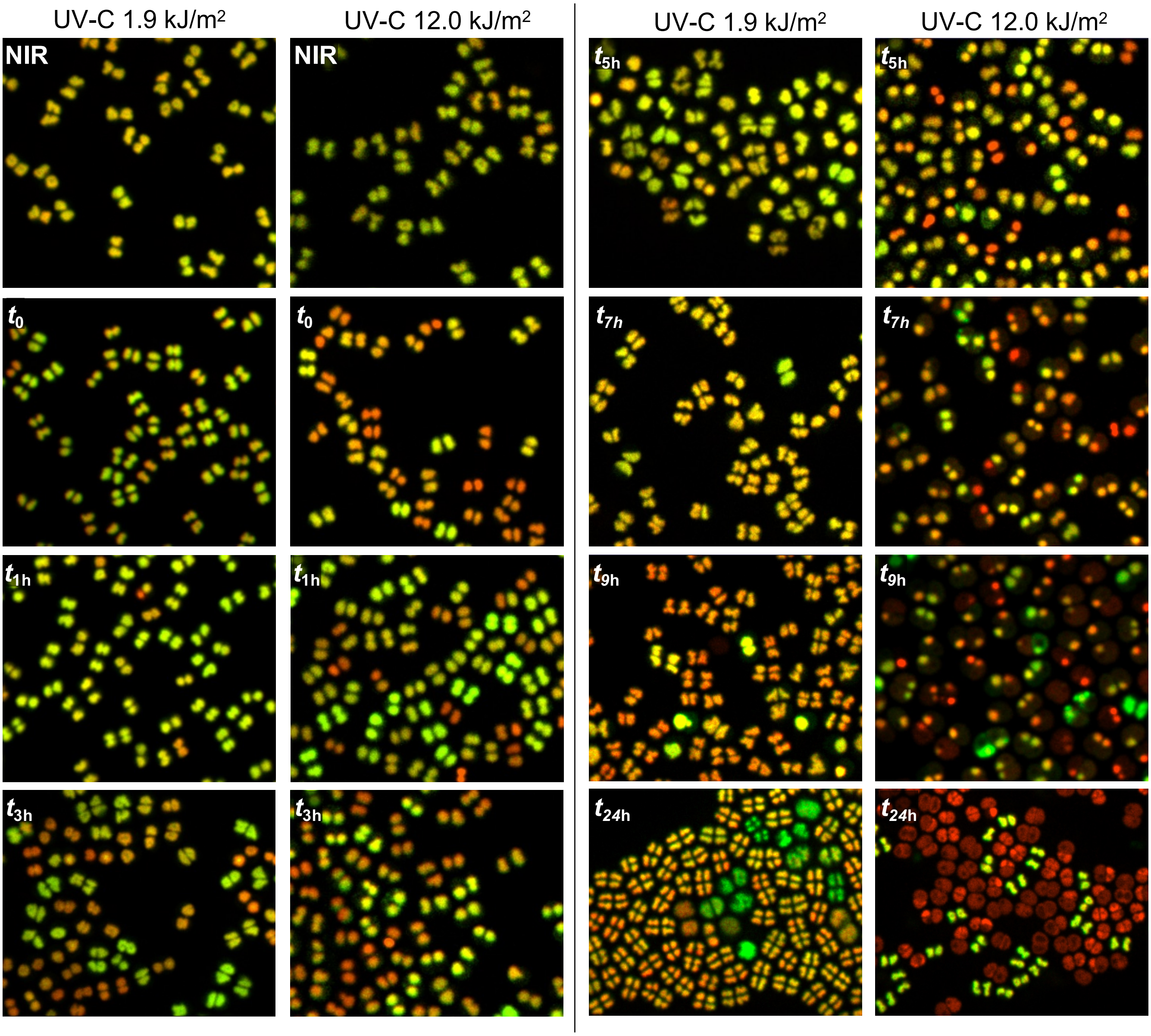


***Figure S14: Representative images of DR$^{HUmCh}$ nucleoids doubly stained with mCherry (red) and Syto9 (green) at different timepoints after exposure to sublethal (1.9 kJ/m2; left column) or lethal (12.0 kJ/m2; right column) UV-C light.*** *NIR: non-irradiated. Yellow cells show similar levels of mCherry and Syto9 staining. Red cells exhibit no or low Syto9 staining as a result of DNA degradation, while green cells are either devoid (as in $t_{24h}$) or express low levels of HU-mCherry (as in $t_{3h}$).*

### Figure S15

**Optimized mEos4b sequence**

gga tcc
atg gtc tcg aaa ggg gag gaa gac aac gtc tcg gcg atc aag ccc gac atg cgg atc aaa
ctc cgc atg gaa ggc aac gtc aac ggc cac cac ttc gtg atc gac ggc gac ggt acg ggc
aag cct tac gag ggc aag cag acg atg gac ctg gaa gtc aag gag ggc ggt cct ctg cct
ttc gcc ttt gac atc ctg acc acg gcg ttc cat tac ggc aac cgg gtc ttc gtg aaa tac
ccg gac aac atc cag gac tac ttc aag cag tcg ttc cct aag ggt tac tcg tgg gaa cgc
agc ctg acc ttc gaa gac ggt ggc att tgc aac gcc cgc aac gac atc acg atg gaa ggt
gac acc ttc tac aac aag gtc cgg ttt tac ggc acc aac ttc ccc gcc aac ggc cct gtc
atg cag aag aag acg ctg aaa tgg gag ccg tcc acc gag aag atg tac gtg cgt gac ggc
gtg ctg acg ggt gac att gag atg gcg ctg ctg ctc gaa ggt aac gcc cac tac cgt tgc
gac ttc cgg acc acg tac aaa gct aag gag aag ggt gtc aag ctg cct ggc gcc cac ttc
gtg gac cac gcc att gag atc ctg agc cac gac aaa gac tac aac aag gtc aag ctg tac
gag cac gcg gtg gcc cac tcg ggc ctg cct gac aac gcc cgg cgt tga tag gca gaa aaa
atc ccc ccg gtg gca atc cgg ggg gtt ttt tac cgg ttg cta cgc ctt cta ga

Additional 5’ sequence: BamHI-N-terminus of mCherry
GGATCCatggtctcgaaaggggaggaagacaac
Additional 3’ sequence: Stop linker-Ter 116sequence-AgeI-linker-XbaI
tgataggcagAAAAAATCCCCCCGGTGGCAATCCGGGGGGTTTTTTACCGGTtgctacgcctTCTAGA

***Figure S15: Sequence details of the mEos4b gene used in* D. radiodurans.**
